# Paranode stability requires UNC5B expression by oligodendrocytes

**DOI:** 10.1101/2020.08.06.234906

**Authors:** Omar de Faria, Diane S. Nakamura, Samuel Clemot, Doyeun Kim, Mihai Victor Mocanu, Roland Pilgram, Jenea M. Bin, Edwin W. Wong, Amir Shmuel, Abbas Sadikot, Susan L. Ackerman, Timothy E. Kennedy

## Abstract

In the mature CNS, netrin-1 is expressed by neurons and oligodendrocytes and implicated in the stability of axo-oligodendroglial paranodal junctions. Here we report that the netrin receptor UNC5B is highly expressed by mature oligodendrocytes and enriched at paranodes. We demonstrate that paranodes become disorganized following conditional deletion of UNC5B in oligodendrocytes, with disruption of the interface between glial loops and detachment of loops from the axon. As a result, Caspr1 and Kv1.1 disperse along the axon, internodes fail to lengthen and compact myelin periodicity is reduced. Paranodal and axoglial domain disorganization progressively worsens and a delay in motor learning develops in aged mice lacking oligodendroglial UNC5B. Altered glial loop ultrastructure and reduced levels of claudin-11 and JAM-C tight junction proteins support the conclusion that disruption of autotypic junctions between paranodal loops underlies paranode disorganization. Our findings reveal an essential contribution of oligodendroglial UNC5B at paranodes that is required for the stability of mature myelin.

## INTRODUCTION

Netrins are a family of laminin-related proteins that guide cell migration and axon extension in the developing nervous system (Kennedy et al., 1994; Serafini et al., 1994). Netrin-regulated motility is mediated by a group of transmembrane receptors, all members of the Immunoglobulin (Ig) superfamily, that include Deleted in Colorectal Cancer (DCC), the DCC paralogue neogenin, and the four mammalian UNC5 homologues, UNC5A-D. Although DCC signaling to direct cytoskeletal rearrangement has been well studied, UNC5 homologue signaling remains relatively poorly understood (Lai Wing Sun et al., 2011). Furthermore, while netrin-1 function is best understood during embryogenesis, netrin-1 and netrin receptors are also expressed during postnatal maturation in the adult brain and spinal cord (Manitt et al., 2001; Manitt et al., 2004). Consistent with this, netrin-1 and DCC regulate several aspects of postnatal development and adult CNS function, including synaptogenesis, synaptic plasticity, memory and the maintenance of myelin domains (Bull et al., 2014; Glasgow et al., 2018; Glasgow et al., 2020a; Glasgow et al., 2020b; Goldman et al., 2013; Horn et al., 2013; Jarjour et al., 2008; Wong et al., 2019).

Distinct axoglial domains - the node, paranode, juxtaparanode, and internode - are assembled along myelinated axons to enable saltatory conduction of the action potential. The paranode, immediately adjacent to the node of Ranvier, is the myelin domain where compact myelin opens into cytoplasm-containing glial loops. Paranode organization relies on adhesive cell-cell junctions, among them septate-like junctions, that mediate adhesion between oligodendroglial loops and the axolemma, specifically between glial Neurofascin-155 and axonal Contactin and Caspr1 (Boyle et al., 2001; Charles et al., 2002; Einheber et al., 1997; Tait et al., 2000). At the oligodendroglial paranodal loop-loop interface, Claudin-11 contributes to form a specialized autotypic tight junction (Gow et al., 1999). The intact paranodal junction functions as a barrier, segregating potassium channels to the juxtaparanode and sodium channels to the node of Ranvier (Pedraza et al., 2001). Mice null for Neurofascin, Contactin or Caspr1 exhibit loss of septate-like junctions and disorganized paranodes, with ion channels not properly segregated and impaired saltatory conduction (Bhat et al., 2001; Boyle et al., 2001; Dupree et al., 1998; Marcus et al., 2006; Schaeren-Wiemers et al., 2004; Sherman et al., 2005).

UNC5 homologue expression increases with maturation in the CNS (Manitt et al., 2004). The extracellular region of UNC5 homologues is composed of two Ig domains, which bind netrin-1, followed by two Thrombospondin type 1 domains (Geisbrecht et al., 2003; Leonardo et al., 1997). The intracellular region of UNC5s includes a ZU5 domain, homologous to the septate-like junction adaptor protein Ankyrin B and the tight junction adaptor protein Zona Occludens-1 (ZO-1) (Schultz et al., 1998). UNC5B (also called UNC5H2) is the most highly expressed UNC5 family member in CNS white matter, suggesting that it may participate in netrin-1 signaling in oligodendrocytes (Manitt et al., 2004). Here, we show that *unc5b* expression is upregulated in the adult mouse brain and that UNC5B protein is enriched at paranodal junctions along myelinated axons. We report that following cre-mediated excision to delete UNC5B from oligodendrocytes *in vivo*, paranode ultrastructure is disrupted, axonal domain segregation is compromised, compact myelin periodicity is reduced, and internodes fail to lengthen with age. Notably, paranodal disruption is less severe in young animals and defects worsen with age. Consistent with this, motor learning is impaired in aged mice that lack oligodendroglial UNC5B. We detected reduced amounts of the paranodal tight junction proteins Claudin-11 and JAM-C in UNC5B cKOs providing evidence that disruption of the autotypic junctions between glial loops underlies paranodal disorganization. Our findings indicate that expression of UNC5B by oligodendrocytes is required to maintain the organization of myelin in the mature CNS.

## RESULTS

### UNC5B is enriched at paranodes of myelinated axons in the adult brain

*Unc5b* is highly expressed in adult rat CNS white matter, suggesting that UNC5B protein might contribute to netrin-1 signaling in mature oligodendroctyes (Golan et al., 2008; Manitt et al., 2004). To investigate UNC5B function in the mature CNS, we first characterized the distribution of *unc5b* expression in the mouse brain. RT-PCR analysis of whole brain cDNA revealed increasing levels of *unc5b* mRNA during the first postnatal month that peak at ~P30, the age corresponding to peak expression of PLP, the most abundant protein in CNS myelin (Cook et al., 1992; Macklin et al., 1991)(Fig. 1A). *Unc5b* expression gradually decreases at subsequent ages, with relatively high levels of mRNA maintained at up to 12 months of age (Fig. 1A; expression at P360 is approximately half the peak expression at P30). UNC5B protein was not detected in lysates of P1 cerebellum, but was abundant in lysates from P60 mice, confirming preferential expression in the mature CNS (Fig. 1B).

**Figure 1:**
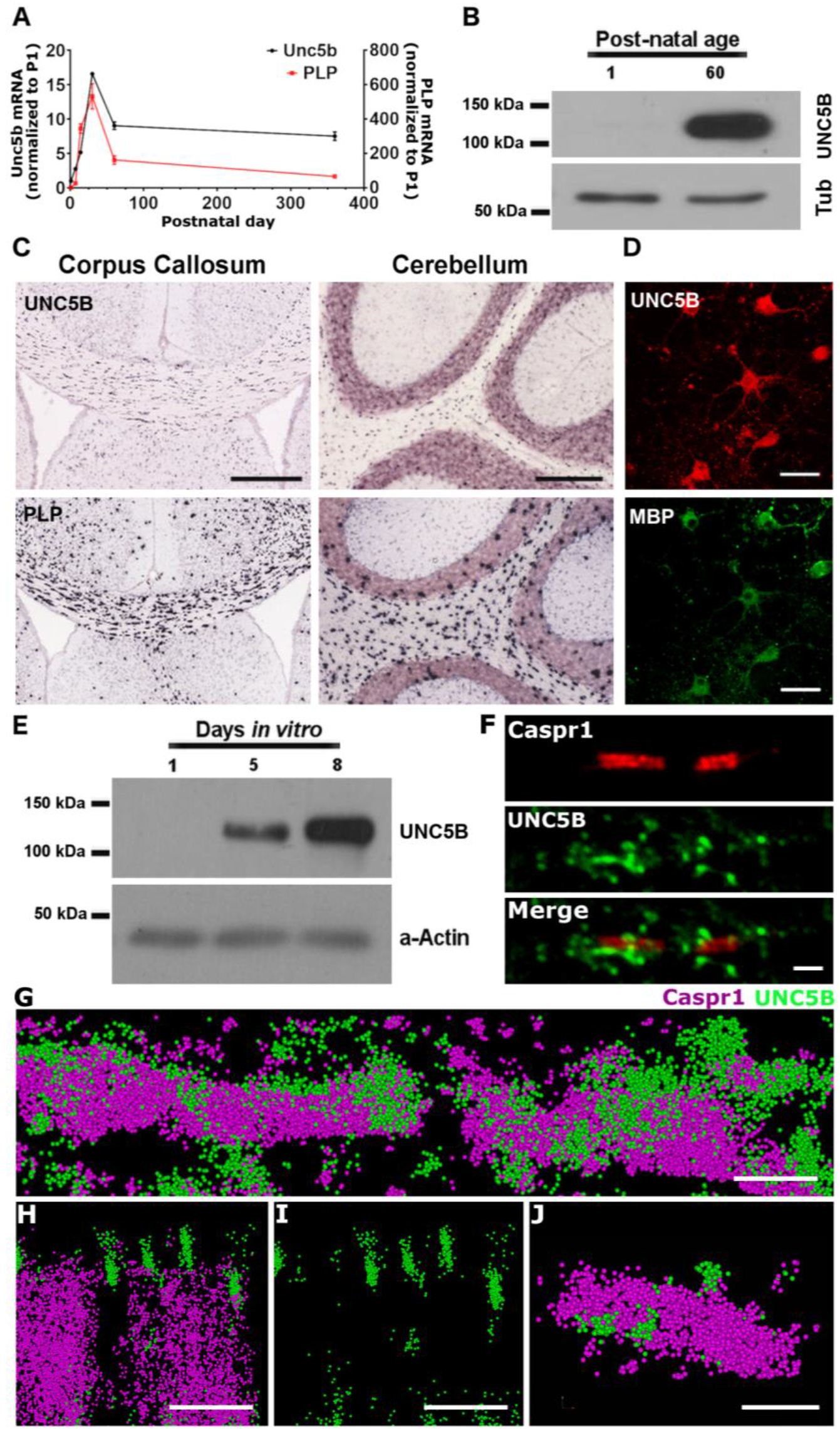
UNC5B is enriched at oligodendroglial paranodal junctions in adult mouse brain. **(A)** Real-time RT-PCR analysis of *unc5b* and *plp* expression in the mouse brain. **(B)** Western blot of cerebellum lysates reveals readily detectable amounts of UNC5B protein in the adult CNS. **(C)** *In situ* hybridization of *unc5b* mRNA indicates expression in white matter tracts of corpus callosum and cerebellum. PLP expression is shown for comparison (adapted from the Allen Mouse Brain Atlas (Lein et al., 2007); (http://mouse.brainmap.org/gene/show/18587 and http://mouse.brain-map.org/experiment/show/71656860). Scale bar: 420 and 210 μm. **(D)** UNC5B and MBP immunostaining of mature oligodendrocytes cultured *in vitro* for 8 days. Scale bar: 32.5 μm. **(E)** Western blot analysis of UNC5B in cultures of purified oligodendrocytes at 1, 5 and 8 DIV. **(F)** Caspr1 and UNC5B immunostaining of myelinated axons teased from adult mouse spinal cord, showing UNC5B and Caspr1 proteins enriched at the paranode. Scale bar: 3 μm. **(G-I)** STORM super-resolution imaging of a section of adult mouse optic nerve revealing UNC5B (green) immunostaining spiraling around the Caspr1 (magenta) positive domains. **H** and **I** correspond to the same paranode. Scale bar: G, H and I: 1 μm; J: 700 nm.

We then examined the distribution of *unc5b* expressing cells in the adult brain (Fig. 1C). Positive *in situ* hybridization was detected in cells throughout the white matter of the corpus callosum and cerebellum, similar to PLP expression in the adult mouse brain (Allen Brain Mouse Atlas; Lein et al., 2007). To confirm that UNC5B protein is expressed in the oligodendrocyte lineage, we cultured oligodendrocyte precursor cells (OPCs) from the brains of newborn rat pups and immunolabelled for UNC5B and the oligodendrocyte marker MBP. Following OPC differentiation, at 8 DIV, UNC5B immunoreactivity was readily detected in MBP^+^ oligodendrocytes (Fig. 1D) and, accordingly, immunoblot analysis revealed increasing levels of UNC5B protein as OPCs differentiate (Fig. 1E). To characterize the distribution of UNC5B along myelinated axons, immunolabelled teased fibers from adult mouse spinal cord were imaged using confocal microscopy. We observed UNC5B partially overlapping with the paranodal protein Caspr1 (Fig. 1F). To further characterize UNC5B distribution at the paranode, we performed Stochastic optical reconstruction microscopy (STORM) super-resolution imaging of immunostained sections of adult mouse optic nerve. STORM revealed UNC5B spiraling around Caspr1, which marks the axonal plasma membrane at the paranode (Fig. 1G). UNC5B localized apart from, but immediately adjacent to Caspr1 (Fig. 1G-J), in a distribution suggesting enrichment at paranodal autotypic loop-loop junctions. Collectively, our findings indicate that *unc5b* is expressed by oligodendrocytes and UNC5B protein is enriched at paranodes along myelinated axons in the adult mouse brain.

### Oligodendrocyte-specific deletion of UNC5B

The enrichment of UNC5B at paranodes suggests a possible contribution to the organization and maintenance of paranodal junctions. *Unc5b* null mice die during early development, before embryonic day 14 (Lu et al., 2004), precluding the use of *unc5b* nulls to study loss of function at later ages. We therefore generated a transgenic mouse line carrying a floxed *unc5b* allele to study UNC5B function *in vivo* (Fig. 2A, B). To selectively delete *unc5b* from oligodendrocytes, mice carrying the floxed *unc5b* allele were crossed with Olig2-Cre expressing mice (Schuller et al., 2008). To confirm the identity of the Cre-expressing cell types, we crossed Olig2-Cre mice with the ROSA26 reporter line (Soriano et al., 1998). β-galactosidase activity assays performed on brain sections of these mice revealed Cre expression by cells in the corpus callosum and cerebellar white matter, consistent with Cre expression by oligodendrocytes (Fig. 2C). Cre is also expressed by motor neurons in the ventral spinal cord of Olig2-Cre mice; however, these cells do not express *unc5b*, but rather *unc5a* and *unc5c* (Burgess et al., 2006; Dillon et al., 2007; Leonardo et al., 1997).

**Figure 2:**
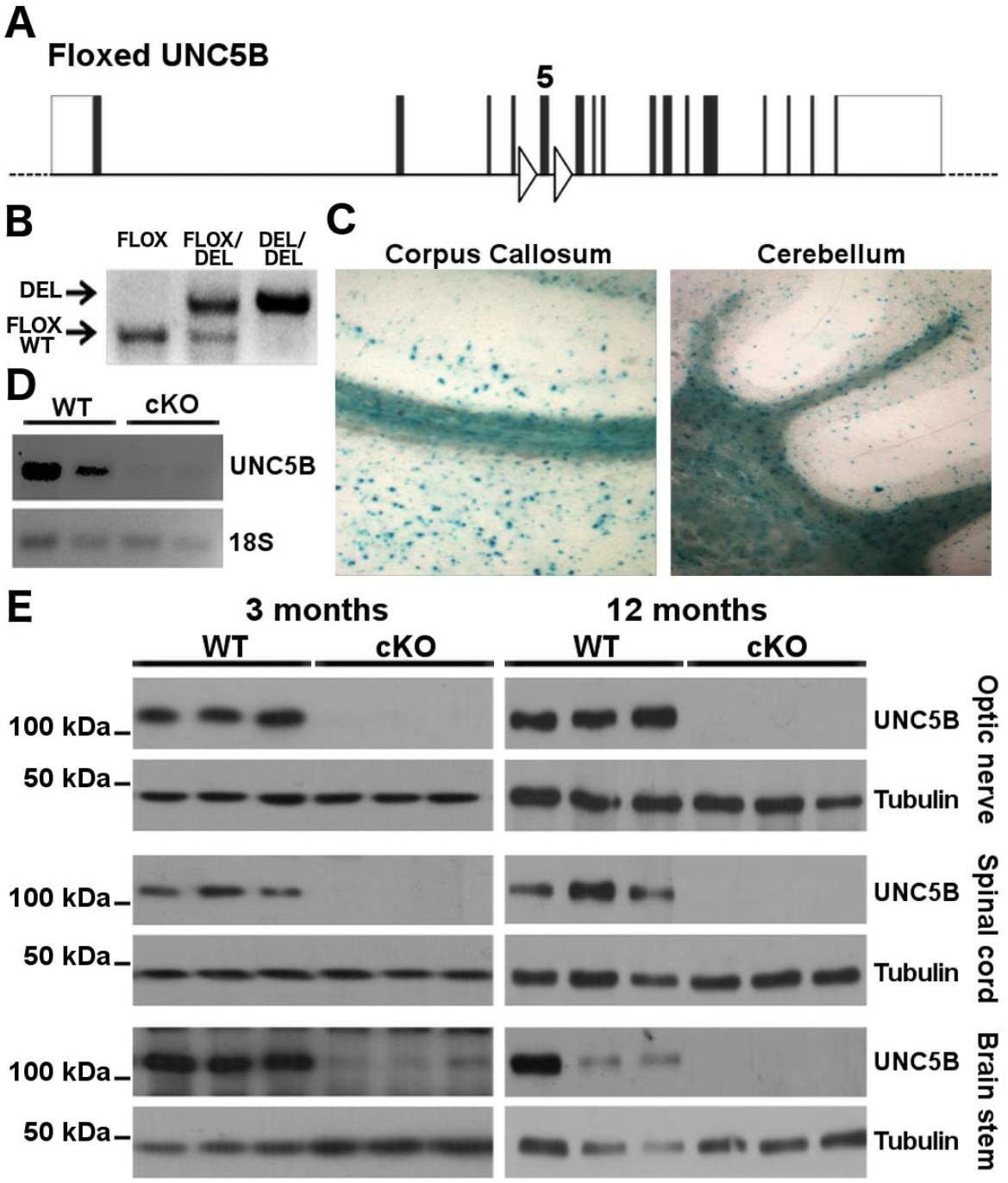
Conditional deletion of *unc5b* expression from oligodendrocytes in UNC5B cKO mice. **(A)** Schematic representation of the floxed *unc5b* allele and **(B)** representative image showing genotyping of *unc5bFlox* mice. **(C)** β-Gal staining of brain sections of adult Olig2-Cre/ROSA26 reporter mice shows Olig2-driven Cre expression in oligodendrocytes of corpus callosum and cerebellum. **(D)** RT-PCR analysis of *unc5b* expression in brain stem lysates of 3 month-old cKO mice shows reduced *unc5b* mRNA expression compared to WT controls. **(E)** Western blot analysis of 3 and 12 month-old mouse optic nerve, spinal cord and brain stem lysates confirms deletion of *unc5b* expression from oligodendrocytes.

To confirm the reduction of *unc5b* expression in Olig2-Cre/*unc5b*flox mice (subsequently referred to as UNC5B cKOs), we carried out RT-PCR analyses of lysates derived from the brain stem of 3 month-old UNC5B cKOs and wild-type littermates. *Unc5b* mRNA was readily detected in *wild-type* tissue, but not in lysates from UNC5B cKO tissue at this age (Fig. 2D). Western blot analyses were then performed on lysates from three white matter-rich regions, optic nerve, spinal cord and brain stem. At 3 and 12 months of age, levels of UNC5B were substantially reduced in all CNS regions analyzed, confirming that *unc5b* is predominantly expressed by oligodendrocytes in the CNS (Fig. 2E). Levels of netrin-1 protein in the white matter of UNC5B cKOs were comparable to wild-type (not shown).

### UNC5B cKO mice display normal myelin abundance but reduced myelin periodicity

To determine whether UNC5B is required for normal CNS myelination, we first analyzed the expression of the major myelin proteins MBP and PLP in 6-9 month-old UNC5B cKO and wild-type mice. Western blot analyses of lysates from white matter-rich CNS regions revealed comparable expression levels of MBP and PLP in UNC5B cKO and wild-type littermates (Fig. 3A). Electron microscope (EM) analyses performed on sections of optic nerve from 7 month-old UNC5B cKOs and wild-type animals revealed no difference in the percentage of axons myelinated at this age (91.12±2.03 for WT and 93.99±1.45 for UNC5B cKO, Fig. 3B, C). In addition, quantification of the g-ratio of myelinated fibers revealed no difference in myelin thickness or axon diameter in 7 month-old UNC5B cKO animals compared to wild-type controls (g-ratio=0.605±0.012 for WT and 0.599±0.013 for UNC5B cKO, Fig. 3D, E). The number of outfoldings of residual compact myelin and the size of the inner tongue were also not affected by UNC5B deletion (Fig. 3J), in contrast to the increased outfoldings of residual myelin detected in mice with DCC deleted from mature oligodendrocytes (Bull et al., 2014).

**Figure 3:**
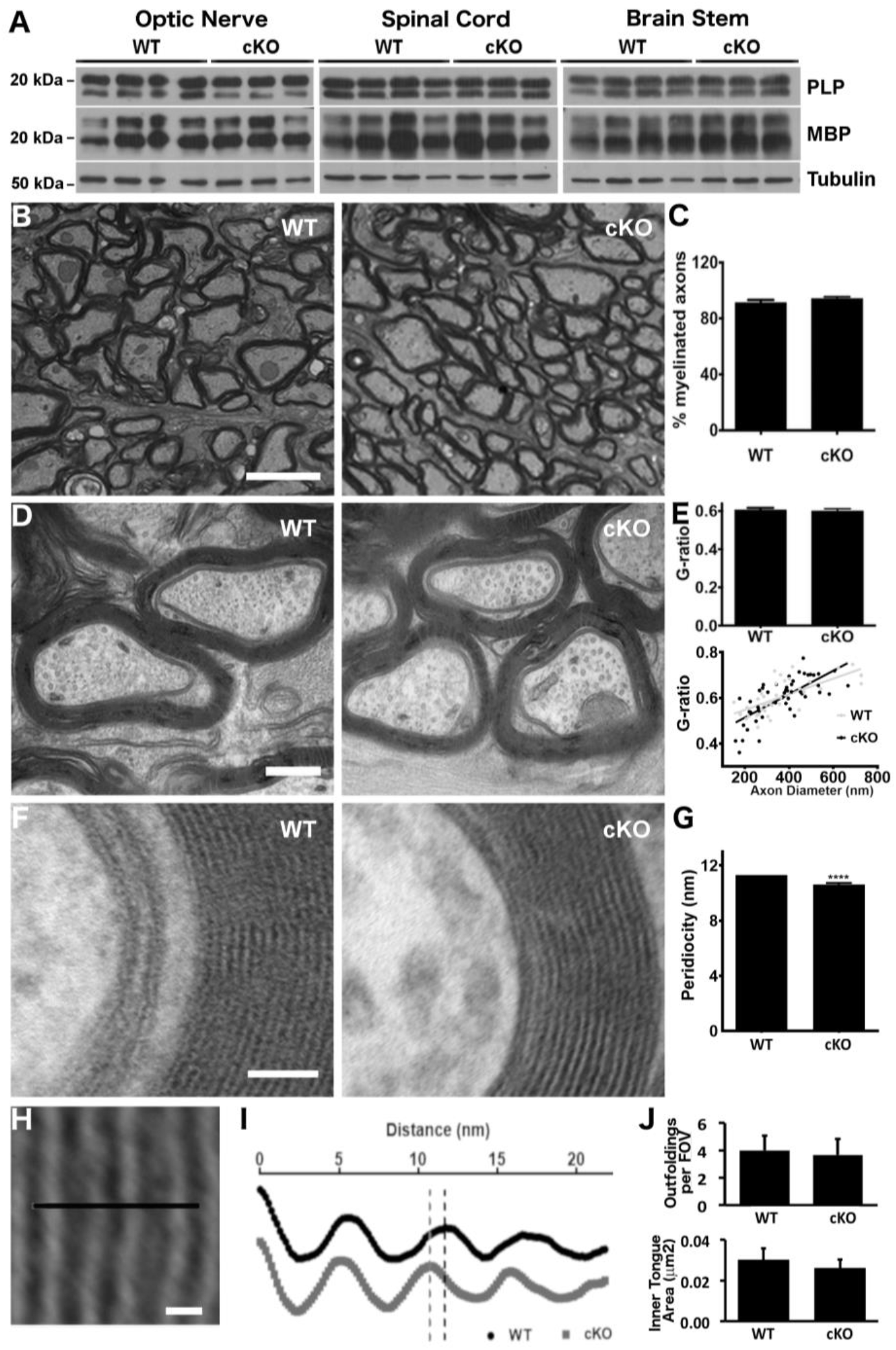
Unchanged myelin abundance with reduced compact myelin periodicity in UNC5B cKO mice. **(A)** Western blot analysis of major myelin proteins PLP and MBP in optic nerve, spinal cord and brain stem lysates of 6-9 month old cKO and wild-type littermates. **(B,C)** Percent of myelinated axons and **(D,E)** g-ratios were measured in electron micrographs of coronal sections of optic nerve from 7 month old UNC5B cKO and wild-type mice. Scale bar: **(B)** – 2 μm **(D)** – 300 nm. 450-650 axons were analyzed to calculate percent of myelinated axons; 40-50 axons were analyzed for g-ratio quantification. **(F)** Representative micrographs of wild-type and cKO optic nerve sections showing general features of compact myelin. Scale bar: 40 nm **(G)** Mean periodicity is reduced by deletion of *unc5b* expression. >100 individual periods were measured for both genotypes. **(H, I)** Intensity profiles across the line at the center of the micrograph (spanning 2 periods) were collected for 20-25 micrographs and averaged. The vertical dotted lines represent the end of a myelin period. Myelin periodicity is ~1 nm shorter in UNC5B cKOs. Scale bar in **(H)**: 10 nm. **(J)** The number of myelin outfoldings and the area of the inner tongue in EM cross-sections of the optic nerve are comparable in UNC5B cKO and wild-type animals. Inner tongue area was analyzed in 20 axons. Bar graphs are plots of means and error bars indicate ±SEM. \*\*\*\**p* < 0.0001 (unpaired student-t test).

We then examined if UNC5B is required for the proper organization of compact myelin. Ultrastructural analysis of adult UNC5B cKO optic nerve identified no apparent defects in compact myelin organization and structure: major dense lines and intraperiod lines appeared normal, the mean number of wraps did not differ from wild-type, and no signs of degeneration were observed (Fig. 3F). However, myelin periodicity, measured from the apex of one major dense line to the next, was significantly, albeit slightly, decreased around UNC5B cKO fibers (Fig. 3G-I). Interestingly, the opposite phenotype, a small, but significant increase in compact myelin periodicity was previously reported in cultured cerebellar organotypic slices derived from mice null for the netrin-1 receptor DCC (Jarjour et al., 2008).

### Paranode ultrastructure is disrupted in UNC5B cKOs

Paranodes were previously reported to be disorganized in long-term, ~60 DIV, but not short-term, ~25 DIV, organotypic cultures derived from netrin-1 null or DCC null newborn mice (Jarjour et al., 2008). *In vivo*, paranodal organization progressively degraded in mice with selective deletion of DCC from mature oligodendrocytes in the adult CNS, confirming the role of oligodendroglial DCC in paranode maintenance (Bull et al., 2014). To determine whether UNC5B also contributes to paranodal organization, we examined paranodal ultrastructure in adult UNC5B cKO mice. EM micrographs of 7 month-old wild-type optic nerve revealed properly aligned glial loops well attached to the axolemma, with no appearance of degeneration. In contrast, 7 month-old UNC5B cKO paranodes were disorganized, with glial loops improperly aligned and everted away from the axon (Fig. 4A). Quantification revealed that the mean number of detached loops per paranode was significantly increased in adult UNC5B cKO animals compared to wild-type littermates (0.32±0.117 for WT and 0.91±0.215 for UNC5B cKO; Fig. 4B). The majority of wild-type paranodes (77%) scored as normal, with no detached loops, in comparison with 39% of UNC5B cKO paranodes, which were more likely to have at least one detached loop (Fig. 4C). No loss of transverse bands (TBs) in UNC5B cKO paranodes was detected (Sup Fig. 1); however, large aberrant vacuolar structures (Fig. 4A, asterisks) and a striking fragmentation of plasma membrane staining at the glial loop-loop interface (Fig. 4A, zoom, Arrowheads; Supplemental Movie 1) were observed in UNC5B cKO but not wild-type paranodes. Further investigating this phenotype, transmission EM tomography detected apparent fractures of closely apposed membranes that extend into the compact myelin, appearing to form channels of discontinuity running through multiple layers of compact myelin (Fig. 4D and E, arrowheads; Supplemental Movie 1). Although fragmented interfaces were occasionally observed in wild-type animals, the mean number of breaks per paranode was substantially increased in 7 month-old UNC5B cKOs (Fig. 4D). This discontinuity of electron dense staining of myelin membranes resembles misplaced radial component, which is typically restricted to the internode (Kosaras and Kirschner, 1990), but aberrantly shifts into the paranode in UNC5B cKOs (Supplemental Movie 1). In summary, our EM analyses identify substantial disruption of paranode ultrastructure in UNC5B cKO mice.

**Figure 4:**
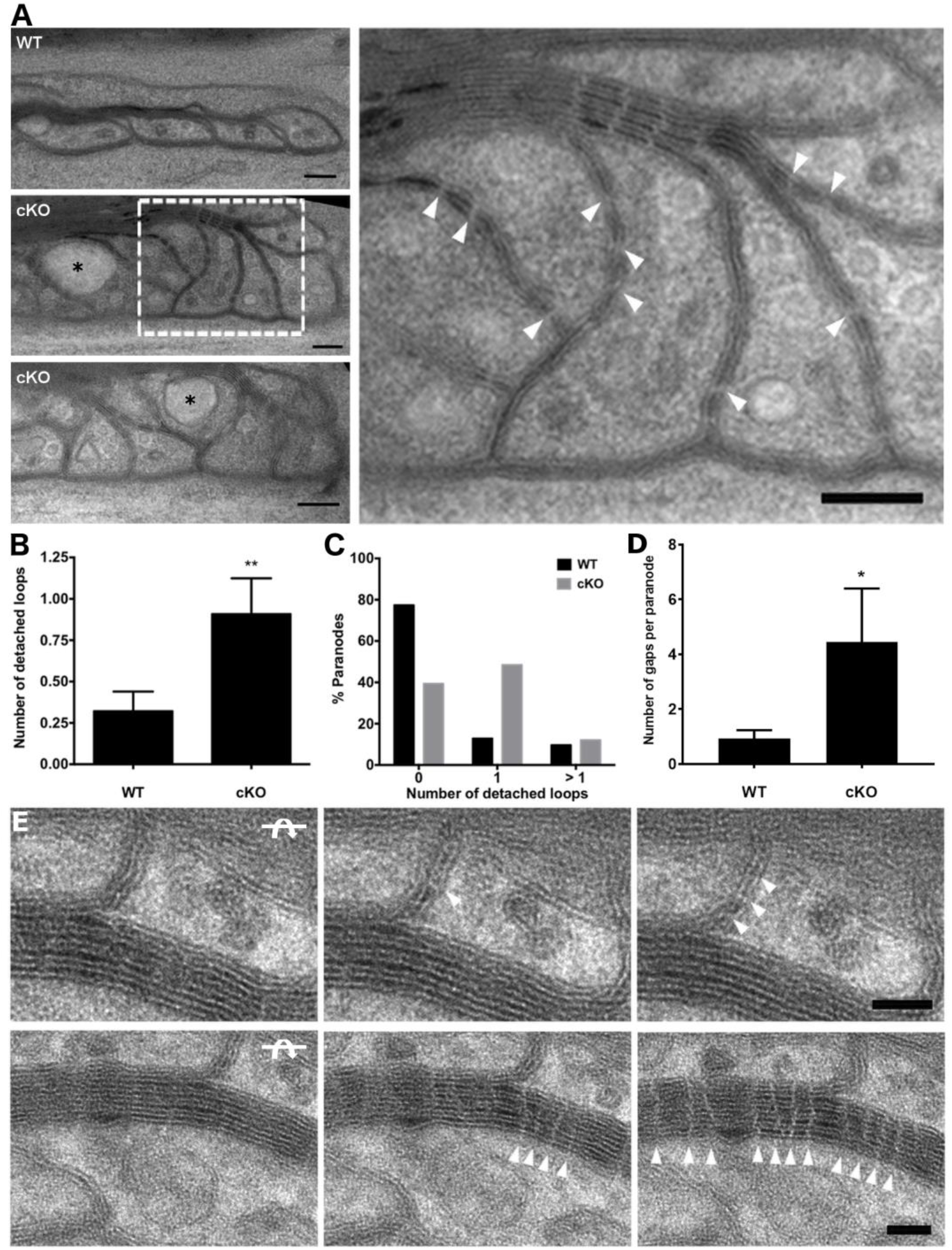
Disruption of paranode ultrastructure in 6-9 month-old UNC5B cKO mice. **(A)** Electron micrographs of longitudinal sections of 7 month-old wild-type and UNC5B cKO optic nerves. Top left panel shows a micrograph of a wild-type paranode and bottom two panels show examples of highly disorganized UNC5B cKO paranodes. The boxed area in the left panel is magnified on the right. Note the segmented loop-loop interface (arrowheads) and large vesicles (asterisks) inside glial loops of cKO paranodes. Scale bars: 100 nm. **(B)** Mean number of detached loops per paranode. ** *p* < 0.01 (Mann-Whitney test). 31 WT and 33 UNC5B cKO paranodes were analyzed. **(C)** Paranode distribution across different levels of disorganization. **(D)** Mean number of gaps detected on glial loop-loop interfaces. * *p* < 0.05 (Mann-Whitney test). 14 wild-type and 19 UNC5B cKO paranodes were analyzed. Bar graphs are plots of means and error bars indicate ±SEM. **(E)** 200 nm longitudinal sections of optic nerve were imaged under a TEM. The microscope stage was tilted by 2-5 degrees between images; tilting progressively reveals fragmentation (white arrowheads) of the glial loop interface (upper panels), which invades the compact myelin (bottom panels). The arrows indicate the direction of tilting of the stage. Scale bar: 100 nm.

### Axoglial domain segregation is disrupted in UNC5B cKO mice

The paranode functions as a molecular “fence” that segregates axonal proteins into distinct domains centered around the node of Ranvier (Pedraza et al., 2001) and disruption of paranodal ultrastructure often results in degradation of myelin domain organization. We therefore investigated if axoglial domains were properly segregated in 6-9 month-old UNC5B cKO mice by immunostaining sections of cerebellum, optic nerve, corpus callosum and deep-layers of neocortex for Caspr1, Kv.1.1 and Na^+^Ch, markers of the paranode, juxtaparanode and node, respectively. To quantify potential defects in domain segregation, we measured the length of these immunopositive domains, the distance between Caspr1 domains flanking a node, and the extent of the overlap between Caspr1 and Na^+^Ch, and between Caspr1 and Kv1.1 (Fig. 5A-C). We detected notable defects in the segregation of axoglial proteins in UNC5B cKOs, with Caspr1 and Kv1.1 domains lengthening in multiple brain regions in comparison to WT littermates (Fig. 5B, C, E, F and Supplementary Fig. 3). In the optic nerve, we detected reduced distance between Caspr1 domains flanking the node of Ranvier and increased overlap between Caspr1 and Na^+^Ch domains at the boundary between paranode and node (Fig. 5F). In some cases, Caspr1 immunoreactivity completely invaded the node of Ranvier and no gap between paranodes was detected, indicating a substantial breakdown of the boundary segregating the paranode and node (Fig. 5C, arrowheads). In contrast, the length of the Na^+^Ch domain and extent of overlap between Caspr1 and Kv1.1 were not different in UNC5B cKO and WTs (Fig 5E, F). Western blot analysis of optic nerve lysates indicated that the loss of domain segregation was not a result of altered levels of Caspr1 expression, which were unchanged in the different genotypes (Fig. 5D). These results indicate that myelin axoglial domains become disorganized following disruption of paranode ultrastructure in 6-9 month-old UNC5B cKOs. As disorganization of the axoglial apparatus is an early hallmark in the autoimmune myelin disorder Multiple Sclerosis (MS) (Dhaunchak et al., 2012; Mathey et al., 2007) we also investigated if perturbation of paranodal organization and axoglial domain segregation might be sufficient to induce an exacerbated immune response in mice lacking oligodendroglial UNC5B. However, examination of microglial cell density in UNC5B cKO and WT animals detected no indication of increased inflammation in UNC5B cKOs (Sup. Fig. 2).

**Figure 5:**
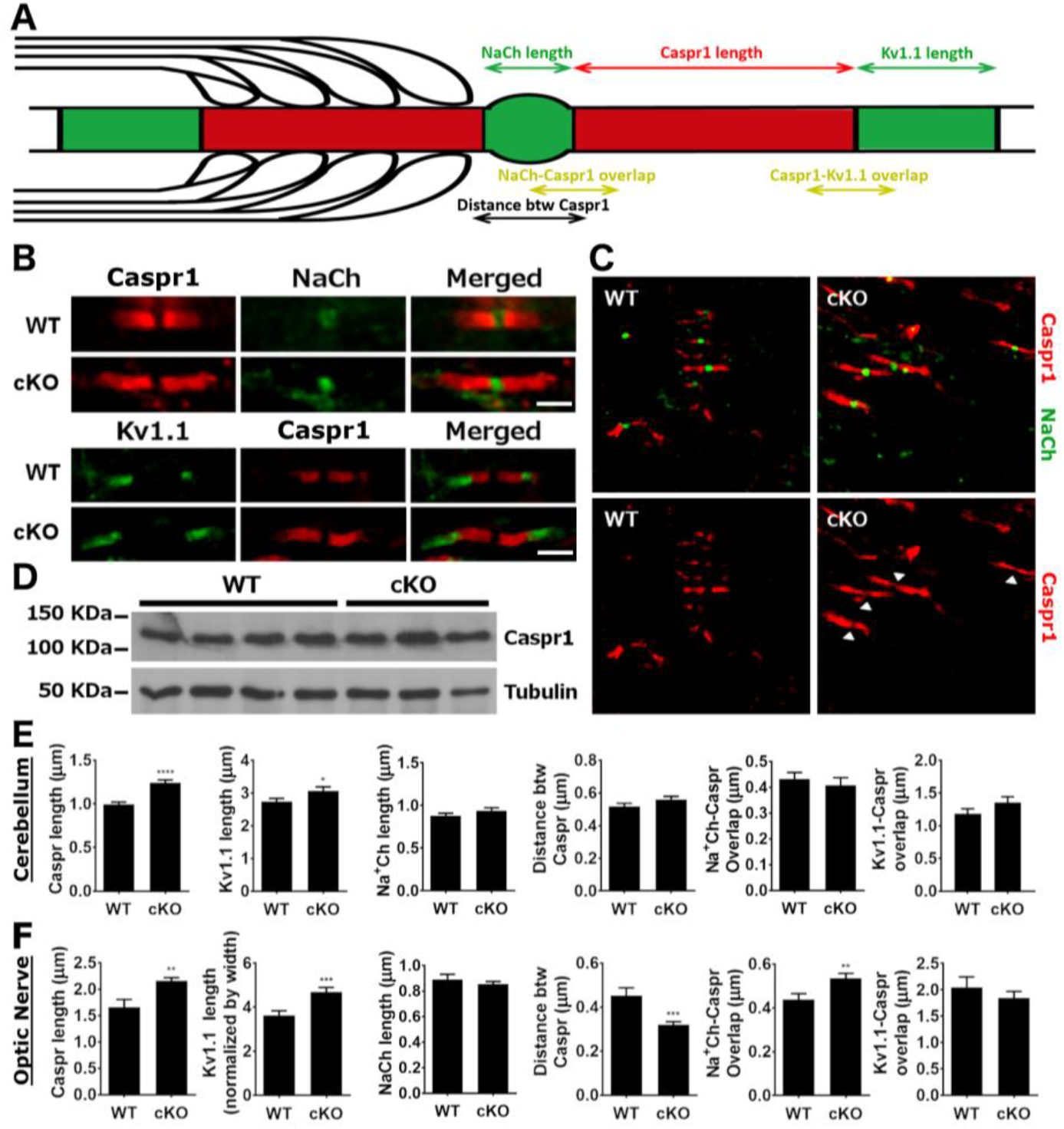
Compromised axonal domain segregation in 6-9 month-old UNC5B cKO mice. Measurement of the length of immunoreactive domains and extent of overlap of Caspr1, Kv1.1, and Na^+^ channel domains along axons in sections of cerebellum and optic nerve of 6-9 month old adult UNC5B cKO and wild-type control mice. **(A)** Schematic representation of the measurements. **(B)** Representative images of Caspr1, Kv1.1 and Na^+^ channel staining in the optic nerve of UNC5B cKO and wild-type mice. **(C)** Caspr1/Na^+^ channel staining of optic nerve showing highly disorganized paranodes in UNC5B cKOs. Arrowheads indicate nodes of Ranvier invaded by Caspr1. **(D)** Western blot analysis of protein lysates from wild-type and UNC5B cKOs reveals unchanged Caspr expression following Unc5b deletion in oligodendrocytes. **(E, F)** Length measurements of immunoreactive domains and extent of overlap between adjacent domains in cerebellum and optic nerve. ** *p* < 0.01; **** *p* < 0.0001 (unpaired student-t test). 150-200 measurements were taken for each immunoreactive domain. Bar graphs are plots of means and error bars indicate ±SEM.

To investigate if changes in the segregation of non-compact myelin domains are accompanied by a corresponding alteration in myelinated internodes, we examined immunostaining in deep-layers of neocortex in sections of UNC5B cKO and WT mice to visualize MBP and Caspr1 (Fig. 6A). Internode length was then quantified as the length of MBP labeled domains flanked by Caspr1 staining. This analysis identified significantly shorter internodal segments in 6-9 month-old UNC5B cKOs compared to age-matched wild-type littermates (Fig. 6B). Taken together, these results reveal degradation of the organization of multiple functional domains along myelinated axons due to the loss of UNC5B function in oligodendrocytes in 6-9 month old cKO mice.

**Figure 6:**
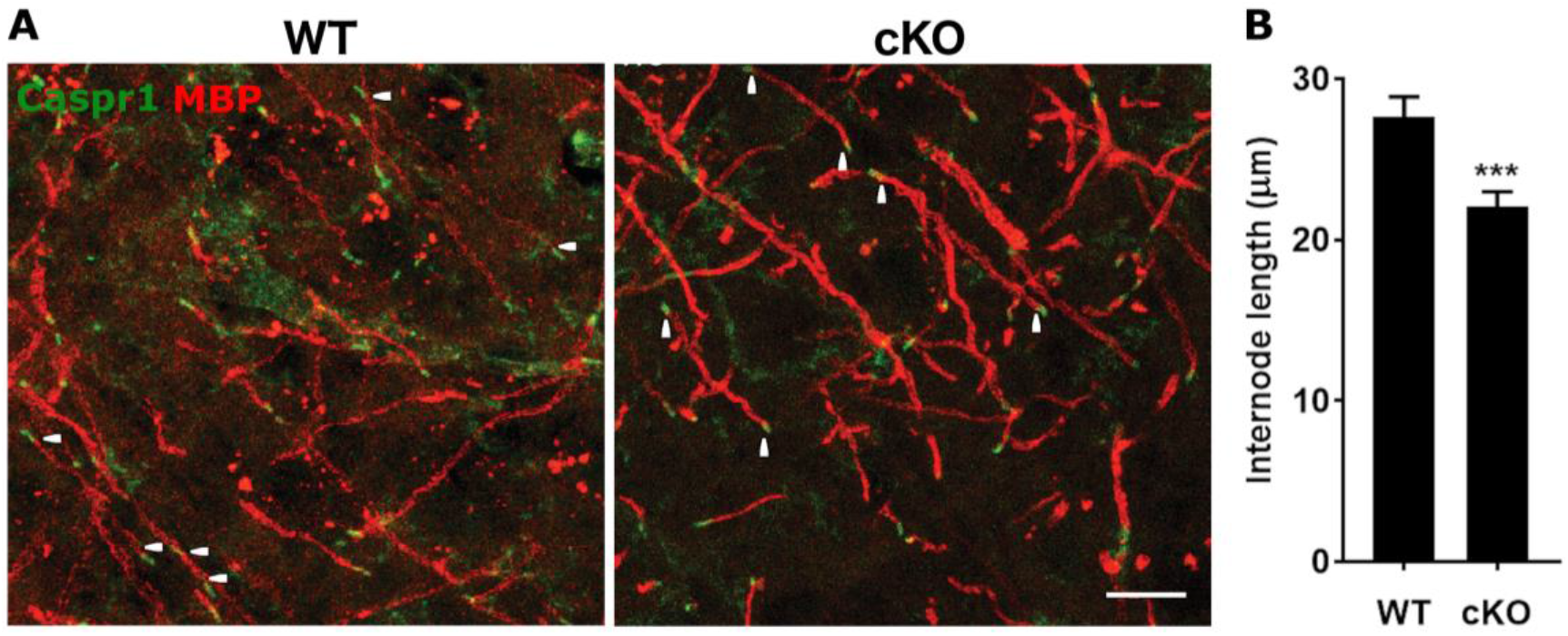
Shorter internodes in 9 month-old UNC5B cKOs. **(A)** MBP and Caspr1 immunostaining in cortical sections from UNC5B cKO and wild-type mice. White arrowheads indicate the end of an internode. Scale bar: 10 μm **(B)** Internode length quantification. ** *p* < 0.001 (Mann-Whitney test) 80 internodes per genotype were measured. Bar graphs are plots of means and error bars indicate ±SEM.

### Paranode disruption increases with age in UNC5B cKOs

To determine if paranodal disruption results from loss of UNC5B function during paranode formation, maintenance or both, we assessed paranode structure in animals at 3 months of age, a developmental stage when the majority of CNS myelination has only recently been completed (Downes and Mullins, 2014; Foran and Peterson, 1992; Hamano et al., 1998). At this age, paranodes in UNC5B cKOs were only slightly more disorganized than those of wild-type age-matched littermates (Fig. 7). EM micrographs revealed that the mean number of detached loops per paranode was marginally increased from 0.8±0.21 in wild-type littermates to 0.93±0.15 in UNC5B cKOs, with 68% of paranodes being scored as normal in wild-types, compared to 39% normal in UNC5B cKO (Fig. 7A-C). This modest increase in paranodal loop disorganization at 3 months of age did not translate into axonal domain disruption, with no increase in paranodal Caspr1 domain length (Fig. 7D-F), and no difference in internode length (Fig. 7G, H). In addition, the average of myelin periodicity, which in 7 month-old UNC5b cKOs is ~1 nm shorter in comparison to WT littermates, was significantly reduced by ~0.5 nm in UNC5B cKOs at 3 months of age (Fig 7I). These results provide evidence that paranodal junctions and internodes in UNC5B cKO mice are more severely disrupted at older ages. While these results do not rule out UNC5B contributing to earlier stages of oligodendrocyte development and myelination, they indicate that UNC5B expression by myelinating oligodendrocytes *in vivo* is essential to maintain paranode and internode organization.

**Figure 7:**
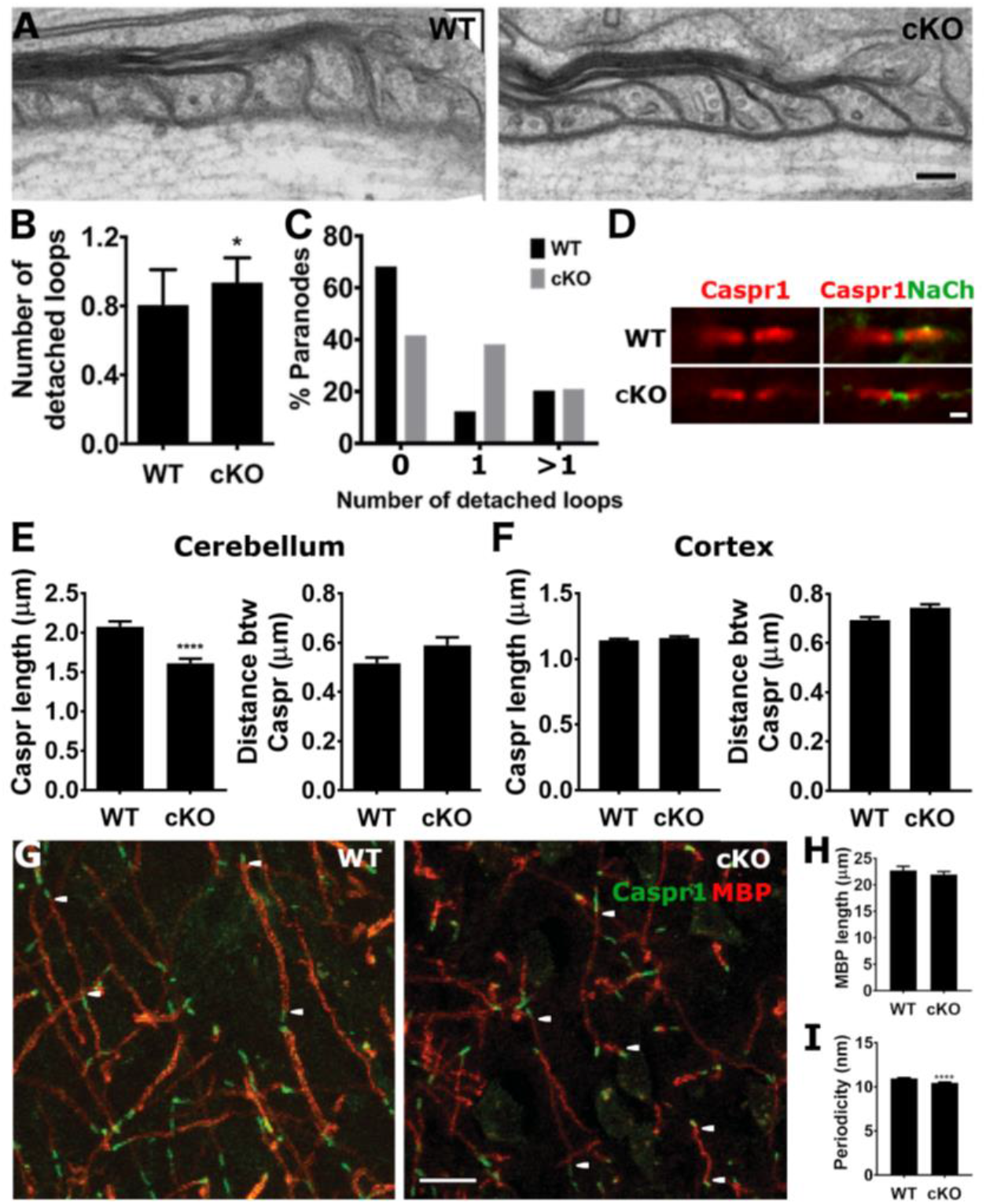
Mild paranode and internode disruption in 3 months-old UNC5B cKOs. **(A)** Representative EM micrographs of optic nerves from 3 month-old UNC5B cKO and wild-type mice. Scale bar: 100 nm. Histograms show **(B)** mean number of detached loops per paranode and **(C)** proportion of paranodes with different levels of disorganization. * *p* < 0.05 (Mann-Whitney test). 50-58 paranodes from 3 animals were analyzed per genotype. **(D)** Representative images of Caspr1 and Na^+^ channel staining in the cerebellum of 3 month-old UNC5B cKO and wild-type mice. Scale bar: 1 μm. **(E, F)** Caspr domain length was measured in the cerebellum and cortex. **** *p* < 0.0001 (unpaired Student’s t-test). **(G)** MBP and Caspr staining on cortical sections from UNC5B cKO and wild-type mice. White arrowheads indicate internode endings. Scale bar: 10 μm. **(H)** MBP domain length. 80 internodes per genotype were measured. **(I)** Myelin periodicity was measured in 32 and 28 axons (~150 periods) from WT and UNC5B cKO animals, respectively. Bar graphs are plots of means and error bars indicate ±SEM.

### Aged UNC5B cKOs develop a motor defect

Mutant mice in which paranodes are severely disrupted typically exhibit some degree of motor defect (Bhat et al., 2001; Boyle et al., 2001; Bull et al., 2014; Dupree et al., 1998; Marcus et al., 2006; Schaeren-Wiemers et al., 2004; Sherman et al., 2005). As UNC5B cKOs did not display overt motor abnormalities, we assessed their behavior for more subtle motor deficits. Animals were examined at 3 months of age, when paranodal disruption is relatively minor, and at 9 and 12 months of age, when disruption is more severe.

Investigating if UNC5B deletion alters spontaneous locomotor activity, we performed an open field analysis. Comparisons between 3, 9 and 12 month-old mice showed a general decrease in locomotor activity in older mice, as expected; however, no differences were found between UNC5B cKOs and wild-type littermates, at any age, in total distance travelled or total active time, supporting the conclusion that spontaneous motor activity is not impaired following UNC5B deletion in oligodendrocytes (Fig. 8A-C). Nor were significant differences detected between UNC5B cKO and WT littermates at any age in the beam walking test, indicating no change in this motivated escape motor response (Kiernan et al., 1999; Wallace et al., 1980) (Sup. Fig. 4). We then investigated if balance and coordination are affected in a context where motor speed and endurance are also challenged, using the accelerating rotarod test. Animals were trained over a period of three days to run on a rotating rod, with the rotational speed increasing every other session of training, and testing on the fourth day. Although the mean latency to fall was not significantly different between genotypes on the test day (100±20.6 s for WTs and 84±10.8 s for UNC5B cKOs; Fig. 8E), we found that during training, 12 month-old but not 3- or 9-month old UNC5B cKOs fell from the rotarod significantly more often than wild-type littermates, specifically on the third (most demanding) day of training (10.8±1.83 for WTs and 16.7±1.82 for UNC5B cKOs, day 3, p < 0.05; Fig. 8D). We conclude that loss of UNC5B from oligodendrocytes results in impaired acquisition of this motor skill with age, as paranode disruption in UNC5B cKO mice worsens over time.

**Figure 8:**
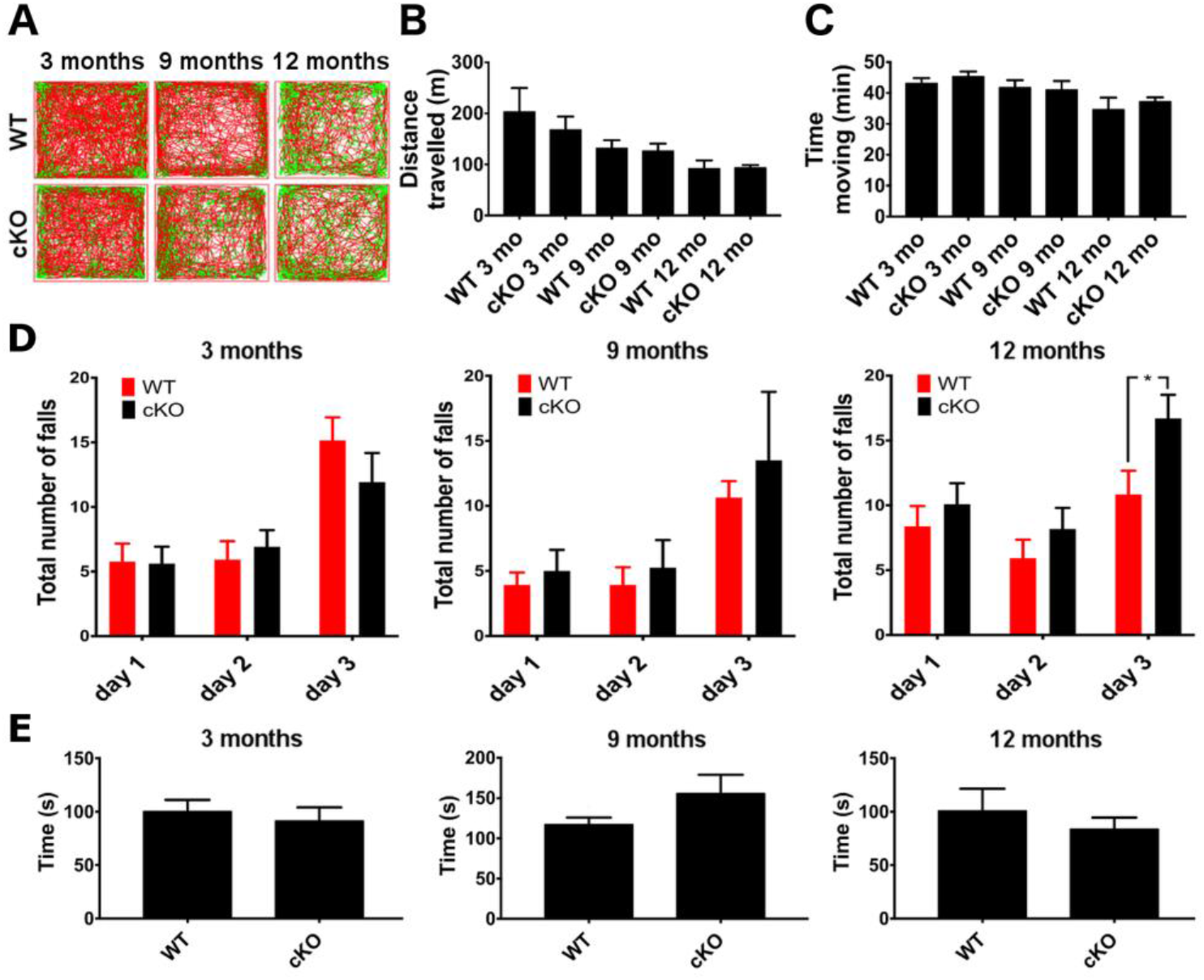
Aged UNC5B cKO mice develop a motor defect. **(A)** Open field analysis of the spontaneous locomotor activity of UNC5B cKO and wild-type littermates. Histograms show **(B)** distance travelled and **(C)** mean active time. n =14 WTs and 13 cKOs at 3 months; 14 WT and 8 cKOs at 6-9 months; 8 WTs and 12 cKOs at 12 months. **(D, E)** Accelerating Rotarod test. Histograms show **(D)** total number of falls on each day of training and **(E)** time mice spent on the rotarod before falling on the test day. n = 13 WTs and 13 cKOs at 3 months; 15 WTs and 8 cKOs at 6-9 months; 12 WTs and 17 cKOs at 12 months. * *p* < 0.05 (Repeated measures 2 way-ANOVA, followed by Sidak’s multiple comparison test). Bar graphs are plots of means and error bars indicate ±SEM.

### UNC5B regulates the expression of paranodal tight junction proteins

The distribution of UNC5B immunoreactivity, organized as a projection perpendicular to Caspr1 marking the axonal surface (Fig. 1G-J), and the apparent fracture of closely apposed glial membranes at paranodes (Fig. 4D) suggested that UNC5B might impact the junctional complex that stabilizes the organization of the interface between glial loops, which, in the CNS, includes the tight junction protein Claudin-11 (Golan et al., 2008; Gow et al., 1999; Poliak et al., 2002). Netrin-1 regulates the expression of tight junction proteins in the polarized epithelial cells that make up the blood-brain barrier (Podjaski et al., 2015), suggesting that tight junctions that link paranodal loops might also be regulated by netrin-1. To test for possible regulation of the glial loop junctional complex by UNC5B, we analyzed tight junction proteins in the optic nerve of 12 month-old UNC5B cKO mice compared to wild-type littermates. UNC5B deletion resulted in reduced Claudin-11 immunofluorescence at paranodes in optic nerve sections (Fig 9A, B), and reduced levels of Claudin-11 and JAM-C proteins in immunoblots of optic nerve lysates (Fig. 9C, D). Together, our findings provide evidence that UNC5B regulates the amount of tight junction protein present at paranodes, and suggest that in the absence of UNC5B reduced levels of these proteins contribute to the disruption of paranodal organization.

**Figure 9:**
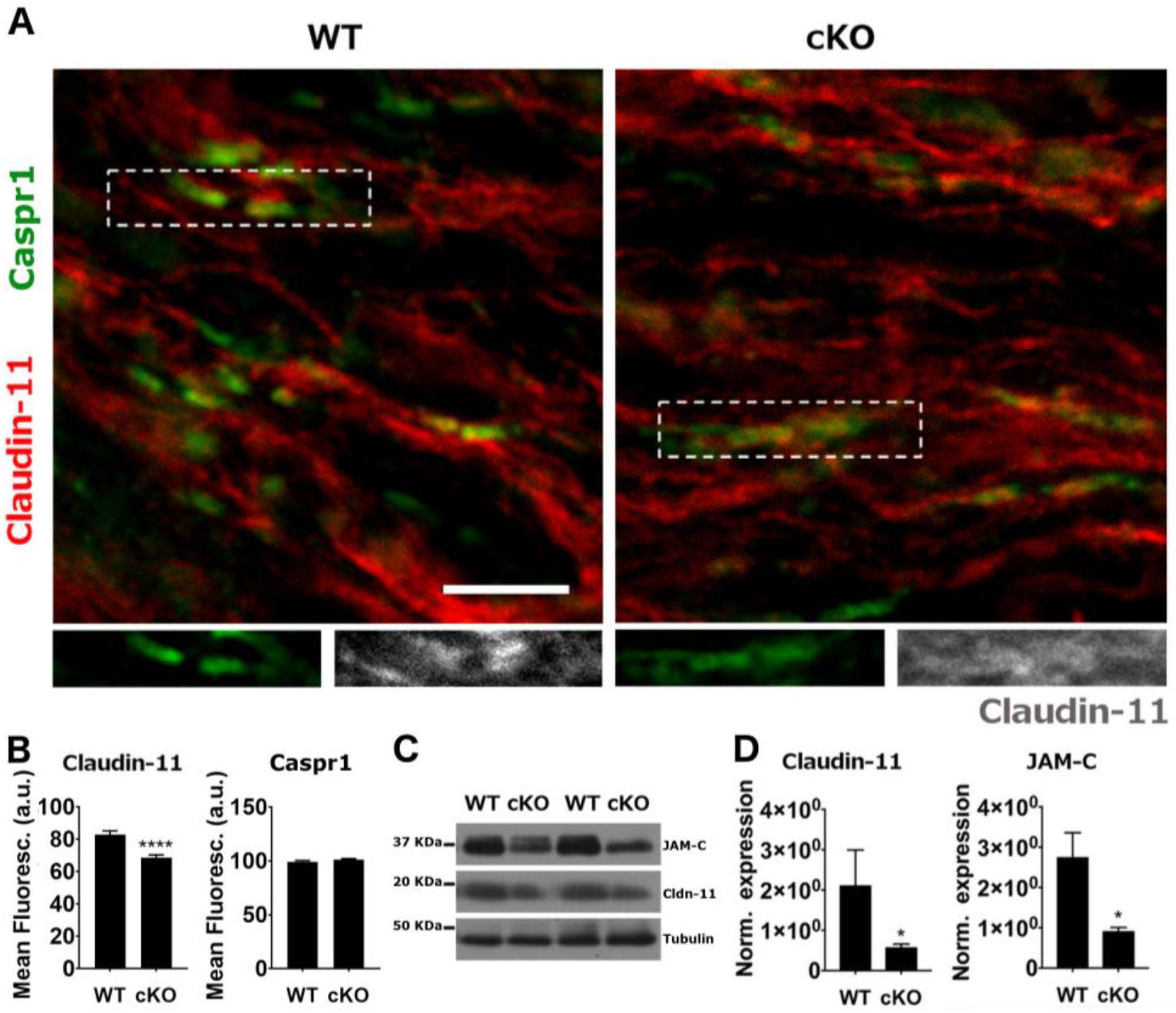
Reduced amounts of Claudin-11 and JAM-C in UNC5B cKO mice. **(A)** Representative images of Claudin-11 and Caspr1 immunostaining in optic nerve sections from 6-9 month old UNC5B cKO and age matched wild-type mice. Scale bar: 5 um **(B)** Quantification of Claudin-11 and Caspr1 fluorescence intensity within the ROI delimited by Caspr1 staining. >300 paranodes were quantified per genotype. **** p < 0.0001 (unpaired Student’s t test) **(C,D)** Western blot analysis of UNC5B cKO and wild-type optic nerve lysates. Graphs show densitometry analysis of JAM-C and Claudin-11 blots. N= 4 animals per genotype. * p < 0.05, ** p < 0.01 (unpaired Student’s t test). Bar graphs are plots of means and error bars indicate ±SEM.

## DISCUSSION

Netrin-1 and its receptors are well known to function in axon guidance in the embryonic CNS. Here, we show that oligodendroglial expression of UNC5B is required to maintain the organization of myelin in adulthood. We found that *unc5b* is highly expressed in the mature CNS and that UNC5B protein is enriched at paranodes along myelinated axons. Oligodendrocyte-specific deletion of *unc5b* expression did not alter the abundance of compact myelin proteins, but resulted in substantial disruption of paranode ultrastructure. As a result, axoglial domain segregation was compromised, with disruption of the distributions of Caspr1 and Kv1.1 along axons, shorter internodes, and reduced compact myelin periodicity. This phenotype was less severe in younger UNC5B cKOs, indicating that the defects worsen with age, ultimately including a motor deficit in aged UNC5B cKOs. We identified reduced levels of Claudin-11 and JAM-C, supporting the conclusion that UNC5B loss disrupts tight junctions at the paranodal glial loop-loop interface. In summary, our results demonstrate that oligodendroglial expression of *unc5b* is required to maintain the organization of mature myelin and axoglial domains in the CNS.

Previously, we reported that in the absence of netrin-1 or DCC, paranodes in organotypic cerebellar slice cultures form properly but become disorganized over time (Jarjour et al., 2008). We then examined a mouse model that selectively deleted DCC from oligodendrocytes in the adult brain and found that paranodes become progressively disordered with age (Bull et al., 2014). These studies provided strong evidence that expression of netrin-1 and DCC by oligodendrocytes maintains the organization of paranodal junctions. In the UNC5B cKO mouse used in here, Cre expression is activated early in postnatal development resulting in Cre recombination prior to completion of myelination (Schuller et al., 2008). Therefore, disruption of paranode organization in 6-9 month-old UNC5B cKOs may result from loss of UNC5B function during paranode formation or maintenance. To investigate this, we examined paranode organization in mice at 3 months of age, when substantial myelination is complete (Downes and Mullins, 2014; Foran and Peterson, 1992; Hamano et al., 1998). We found that paranodes of 3 month-old UNC5B cKOs appear relatively normal, with only a minor subset exhibiting modest paranode disorganization. This suggests an age-dependent phenotype that is consistent with the relatively high levels of *unc5b* transcripts in the brain of mature mice (Fig. 1A). While the modest level of paranodal disorganization at 3 months suggests a possible role for UNC5B in paranode formation, alternatively this may reflect early stages of disruption of previously formed paranodes. Overall, our findings support the conclusion that oligodendroglial UNC5B is essential for paranode maintenance in the mature CNS.

Both UNC5B and DCC are expressed by mature oligodendrocytes, enriched at paranodal junctions and are essential to maintain the organization of axoglial myelin domains, paranodes, and internodes. Although superficially similar, clear differences were detected in the phenotypes of oligodendroglial UNC5B and DCC cKO mice. DCC deletion from mature myelinating oligodendrocytes results primarily in deficits in paranodal axo-glial septate-like junctions (Bull et al 2014), while UNC5B cKOs exhibit disruption of paranodal autotypic loop-loop junctions without loss of axo-glial transverse bands. A dramatic increase in out-folded loops of residual myelin was detected in DCC cKOs, along with an increase in the mean distance between major dense lines in compact myelin, suggesting that loss of DCC results in loosening of compact myelin (Bull et al 2014). No change in residual myelin loops was detected in UNC5B cKOs, and, in contrast to DCC cKOs, UNC5B cKOs exhibit a reduced distance between major dense lines in compact myelin, suggesting tightening of the myelin wrap along the internode. In UNC5B cKOS, but not DCC cKOs, ultrastructural analyses detected striking discontinuities of myelin membranes that resemble radial component, an aligned band of tight junctions that is typically restricted to the internode but aberrantly shifts into the paranode in UNC5B cKOs. We also measured reduced average length of internodes in aged UNC5B cKO mice. This does not appear to reflect internode retraction with aging, but rather that internodes in UNC5B cKOs fail to lengthen normally with age. These findings suggest that loss of UNC5B disrupts the capacity of the internode to be modified during aging, essentially locking-in the juvenile internode length. Although further investigation is required, we speculate that this may contribute to the deficit in motor learning detected in aged UNC5B cKOs.

Our findings suggest that DCC contributes to maintaining appropriate axo-glial junctions, while UNC5B is required for the organization of paranodal autotypic loop-loop junctions. Yet, precisely how DCC and UNC5B interact at paranodes remains to be determined. Studies of migrating cells or axonal growth cones responding to netrin-1 indicate that UNC5 homologues may function with DCC in cis as a heteromeric receptor complex (Lai Wing Sun et al., 2011) or signal independently of DCC (Hong et al., 1999; Keleman and Dickson, 2001). In a study examining blood vessel formation in *Drosophila*, mechanisms that promote cell adhesion and repulsion were found to be spatially segregated to different plasma membrane domains of the same cell, with UNC5 mediating repulsion at lateral domains and DCC adhesion at apical and basal cardioblast contact sites (Albrecht et al., 2011; Macabenta et al., 2013). Our findings suggest that UNC5B and DCC may be similarly spatially and functionally segregated within paranodal loops.

The paranodal autotypic loop-loop interface contains junctional specializations, originally described in epithelial cells, that include tight and adherens junctions (Fannon et al., 1995; Gow et al., 1999). These junctions function as a barrier that prevents the diffusion of small molecules through paracellular spaces and underlie the polarization of epithelial membranes into distinct molecular domains. In the PNS, tight junctions between glial loops contain Claudins-1 and 9 and JAM-C (Scheiermann et al., 2007) whereas Claudin-11 and Connexin-32 are present in tight and gap junctions at CNS paranodes (Gow et al., 1999; Kamasawa et al., 2005; Miyamoto et al., 2005; Poliak et al., 2002). E-cadherin forms adherens junctions between PNS glial loops, whereas adherens junctions have not been detected at CNS paranodes thus far (Fannon et al., 1995). Adaptor proteins, such as Zona Occludens-1, 2 and 3 (ZO-1, 2, 3), link transmembrane junction proteins to the underlying cytoskeleton (Stevenson and Keon, 1998). ZU5 domains are found in the UNC5B intracellular domain and in ZO-1, in which it contributes to linking tight and adherens junctions to the cytoskeleton (Heinemann and Schuetz, 2019; Schultz et al., 1998). Notably, netrin-1 made by vascular endothelial cells regulates tight junction protein levels to control blood-brain barrier permeability (Podjaski et al., 2015), suggesting that paranodal tight junctions may similarly be modulated by netrin-1 signaling. Reduced levels of Claudin-11 and JAM-C proteins in UNC5B cKOs suggest that UNC5B-mediated netrin-1 signaling regulates tight junctions at the glial loop-loop interface and that disruption or mis-positioning of these junctions may underlie paranode disorganization. Claudin-11 null mice lack tight junction intramembranous strands, and although abnormal myelin periodicity or paranode disruption were not reported, aged animals were not examined in these studies (Gow et al., 1999; Maheras et al., 2018). PNS specific deletion of JAM-C, however, results in disorganized Schwann cell paranodes (Scheiermann et al., 2007). Our findings, with these previous studies, suggest that UNC5B interacts with an adhesive complex at the paranodal glial loop-loop interface, and thereby contribute to the maintenance of appropriate junctional specializations.

Examining the ultrastructure of the UNC5B cKO autotypic glial loop-loop interface, we detected crisp regularly spaced discontinuities of electron density in paranodal plasma membrane staining (Fig. 4). High-resolution EM tomography revealed fragmentation extending from the paranode into compact myelin (Fig. 4D). This strongly resembles the morphology of internodal radial component. which corresponds to a band of aligned tight junction-like adhesions that link multiple layers of plasma membrane within compact myelin (Gow et al., 1999; Kosaras and Kirschner, 1990). Our findings suggest that in the absence of UNC5B, with reduced levels of Claudin-11 and JAMC (Fig. 9D), radial component inappropriately invades the paranode. We speculate that this inappropriate positioning of tight junctions may inhibit age related elongation of the internode by compromising the capacity of paranodal loops to reorganize.

No abnormal phenotype was detected in the Open Field test, consistent with the absence of an overt motor defect, and supporting the conclusion that UNC5B deletion does not severely impair behavior. An accelerating rotarod test revealed that UNC5B cKOs were able to learn the task to a level equal to controls, but intriguingly, aged animals exhibited a deficit in the rate of acquisition to learn the motor skill. The delay in motor learning may result from increased fatigue, as myelin disruption can increase fatigue (Simpson et al., 2016), and fatigue is a prominent symptom in demyelinating diseases like multiple sclerosis (Freal et al., 1984; Krupp et al., 1988; Newland et al., 2016); however, UNC5B deletion may also disrupt some aspect of myelin plasticity, which is increasingly understood to play roles in learning, including motor learning (Chorghay et al., 2018; de Faria et al., 2018).

The elucidation of mechanisms that promote myelin maintenance in the adult CNS is critical to the development of new strategies to effectively treat neurological disorders. Multiple Sclerosis (MS), Leukodystrophies, Autism Spectrum Disorder and Schizophrenia are associated with loss of myelin stability and white-matter degeneration (Fields, 2008). Particularly in MS, disorganization of paranodes appears relevant to early disease pathology, preceding disorganization of compact myelin (Howell et al., 2006; Wolswijk and Balesar, 2003). In agreement with these findings, analysis of cerebrospinal fluid samples obtained from children during initial presentation of CNS inflammation did not detect enrichment of proteins associated with compact myelin, but instead identified proteins associated with the node of Ranvier and paranodal junctions, including glial Neurofascin-155 and DCC, enriched in the CSF of children subsequently diagnosed with MS (Dhaunchak et al., 2012). Furthermore, a proteomic screen of human myelin identified Neurofascin-155 as a humoral immune response target in MS (Mathey et al., 2007). These findings indicate that paranodal junctions may be selectively vulnerable to inflammatory attack and suggest that disruption of paranodal organization may be a key event at MS onset. A better understanding of the principles governing paranode formation and maintenance will provide key insights into the dysfunction associated with demyelinating diseases and aid the development of more effective therapies. In summary, our findings reveal an essential contribution of UNC5B in oligodendrocytes to the axoglial apparatus that is required for the maintenance of healthy myelin.

## MATERIALS and METHODS

### Animals

Generation of the targeted *unc5b* allele was performed by homologous recombination in R1 ES cells and targeted cells were identified by Southern blot hybridization. ES cells were injected into B6J blastocysts to generate *unc5b*^neo/+^ mice. *Unc5b*^neo/+^ mice were mated to B6.129S4-*Gt(ROSA)26Sor*^tm1(FLP1)Dym^/RainJ mice (The Jackson Laboratory, stock #009086), to remove the neo cassette. Mice were backcrossed to C57BL6/J mice for ten generations prior to these studies. Olig2-Cre mice (Schuller et al., 2008) were obtained from the Jackson Laboratory. ROSA^26^-LacZ reporter mice (Soriano et al., 1998) were provided by Dr. Jean-François Cloutier (McGill University). All procedures were performed in accordance with the Canadian Council on Animal Care guidelines for use of animals in research.

### Histochemistry, Immunohistochemistry, Immunocytochemistry, and Western blot analyses

The following antibodies were used in this study: mouse monoclonal anti-α-actin (Sigma), mouse monoclonal anti-Caspr1 for immunohistochemistry (UCDavis/NIH NeuroMab Facility), rabbit anti-Kv1.1 (Abcam), mouse anti-pan sodium channel (Sigma), mouse monoclonal anti β-tubulin III (Abcam), goat polyclonal anti-UNC5B for immunostaining (R&D Research), rabbit monoclonal anti-UNC5B for western blot (Cell Signaling), rabbit monoclonal anti-Claudin-11 (Abcam) and rabbit monoclonal anti-JAM-C (Abcam). Rabbit polyclonal anti-Caspr1 (for western blot and immunostaining), rabbit polyclonal anti-CNP, rabbit polyclonal anti-MAG, rabbit polyclonal anti-MBP and mouse monoclonal anti-PLP were gifts from Dr. David Colman (McGill University).

Mice were deeply anesthetized with 4X Avertin and transcardially perfused with phosphate buffered saline, pH 7.4 (PBS) followed by 4% paraformaldehyde (PFA) in PBS. The mouse brain was dissected and post fixed in 4% PFA for 1 hr at 4°C with gentle shaking. Fixed tissue was equilibrated in 30% sucrose, PBS for 1 week at 4°C. After embedding in optimal cutting temperature compound (Sakura Finetek), tissue was cut in 20 μm-thick sections using a Leica cryostat. Slides were air dried for 15 min before being stored at −80°C. For immunohistochemical analyses, slides were allowed to equilibrate to RT and washed 3 times with PBS for 5 min each wash. Sections were blocked at RT with 5% bovine serum albumin (BSA), 0.3% Triton-X, in PBS for 1 h followed by an overnight incubation with primary antibodies at 4°C. Primary antibodies were diluted in 3% BSA, 0.3% Triton-X, in PBS. Sections were then washed with PBS three times, 10 min per wash, and incubated overnight at 4°C with the appropriate secondary antibodies diluted in 3% BSA in PBS. Sections were washed three times with PBS (10 minutes per wash) and briefly dipped into water before mounting. Images were collected using an Olympus Fluoview confocal microscope, Plan Apo 60X 1.42 n.a. oil immersion objective lens, 4X zoom, and a Leica SP8 confocal microscope, 63X, 1.40 n.a. oil immersion objective lens. Measures of length and width of immunoreactive domains were carried out using ImageJ software (Schneider et al., 2012).

For western blot analyses, protein lysates were prepared from freshly isolated mouse optic nerve, brain stem, cerebellum and spinal cord. Isolated tissues were homogenized on ice with RIPA buffer containing protease and phosphatase inhibitors. Equal amounts of protein were resolved by SDS-PAGE and transferred to PVDF membrane (BioRad). Membranes were blocked with 5% milk or 5% BSA in TBS containing 0.1% Tween (TBST) for 1h at RT and incubated overnight with primary antibodies diluted in 1% milk, TBST, at 4°C. Membranes were washed three times with TBST and incubated with horseradish peroxidase-conjugated secondary antibodies for 1h at RT. Membranes were washed and developed with an Enhanced Chemoluminescence Detection kit (Pierce).

For X-Gal histochemical staining, Olig2-Cre/ROSA26 mice were perfused transcardially as described above and 20 μm-thick sections stained for β-galactosidase activity as described (Mombaerts et al., 1996).

### Teased Spinal Cord Fibers

Myelinated axons from the adult spinal cord were isolated for immunohistochemical analysis as described (Jarjour et al., 2020). Adult mice were deeply anaesthetised and perfused transcardially with 4% PFA in PBS. Spinal cords were collected and post-fixed in 4% PFA for 30 min. Meninges were carefully peeled off and white matter regions of the lumber spinal cord microdissected. Isolated white matter tissue was cut into ~2 mm x 1 mm pieces. Acupuncture needles were used to gently tease apart and smear individual myelinated nerve fibers onto SuperFrost Plus slides (Thermo Fisher Scientific, MA, USA).

### Stochastic Optical Reconstruction Microscopy (STORM)

For STORM super resolution microscopy, a wild-type mouse was perfused with 4% PFA. The optic nerve was collected, post-fixed in 30% sucrose and fixed in OCT. 30 μm sections of optic nerve were immunolabeled using the following antibodies: mouse monoclonal anti-Caspr (UCDavis/NIH NeuroMab Facility, CA, USA); goat polyclonal anti-UNC5B (R&D Systems, MN, USA); Alexa 555-conjugated donkey anti-goat; and Alexa 647-conjugated donkey anti-mouse (Invitrogen, CA, USA). Images were taken using an SR-350 Vutara microsope and d-STORM imaging buffer: 1M cysteamine, 2-mercaptoethanol, glucose oxidase, catalase, 1M Tris (pH 8.0), sodium chloride, D-glucose. Z-stacks were acquired in 0.1 μm steps (minimum 25,000 frames per image).

### Electron Microscopy

Mice were deeply anesthetized with 4X Avertin and transcardially perfused with cold 100 mM PBS pH 7.4, followed by cold 2.5% glutaraldehyde, 2% PFA in PBS. Optic nerves were dissected and post-fixed in the same solution for 72 hrs before being processed for electron microscopy, as described (Bull et al., 2014). Images were taken on an FEI Tecnai 12 transmission electron microscope at 120 Kv.

### Behavioral analyses

*Open field test* – mice were allowed to freely explore a 50 x 50 cm arena for 1 h. Locomotor activity was video recorded and tracked using an infrared detector (ViewPoint Life Sciences). VideoTrack software (ViewPoint Life Sciences) was used to analyze motor activity.

*Beam walking test* – mice were trained for four consecutive days, with four sessions per day, to cross a 100 cm-long, 2 cm-wide round beam, elevated 50 cm above the support base, in order to escape to an enclosed box. By day four of training, all animals had learned the task. Because mice wander for variable amounts of time before beginning the task, we determined the 25 cm mark as the task starting point. On the test day, the 2 cm-wide beam used during the training period was replaced by a 1 cm-wide beam. Time to cross the 75 cm distance was measured over four sessions.

*Accelerating Rotarod test* – mice were trained for three consecutive days, with 2-3 sessions per day, to balance on a rotating rod accelerating at 1 revolution/10 seconds (Rotamex, 3 cm diameter rod). The number of falls in each session was scored and pooled by day. On the test day (day 4), acceleration was increased to 1 revolution/8 seconds and latency to fall was recorded over six sessions.

### Real-time RT-PCR

First-strand cDNA synthesis was performed as described (de Faria et al., 2012). Briefly, 50 ng of total RNA was added to 300 ng of random hexamer primers and incubated at 65°C for 10 min. Next, a master mix containing 1X reverse-transcription buffer, 2 U/μl ribonuclease inhibitor (Invitrogen), 1 mM dNTP, and 10 U/μl reverse transcriptase (Invitrogen) was added. RT reaction parameters were as follows: 25°C for 5 min, 50°C for 30 min, 55°C for 30 min, 70°C for 15 min. For real-time RT-PCR, 20 ng of cDNA was incubated with 5 μM forward and reverse primers and 2X SYBR Green PCR master mix (Invitrogen, catalogue #4364344). Reactions were performed using an Applied Biosystems 7000 thermocycler. Data were analyzed using the 2^-[delta][delta]Ct^ method (Schmittgen et al., 2000; Winer et al., 1999) and final expression values normalized to 18S RNA levels.

## Statistical Analysis

Data were analyzed using GraphPad Prism software. Significant differences are indicated with *p*-values: \**p*<0.05, \*\**p*<0.01 and \*\*\**p*<0.001. Statistical tests used are indicated in the figure legends.

## ACKNOWLEDGMENTS

We thank Zahraa Chorghay, Andrew Jarjour and members of the Kennedy lab for comments on the manuscript. This work was supported by operating grants from the Multiple Sclerosis Society of Canada, the Canadian Institutes of Health Research, and the National Institutes of Health (NINDS). OFJ was supported by a Waugh Family MS Society of Canada PhD studentship and TEK was supported by a Scholarship from the Killam Trust and by a Chercheur Nationaux Award from the Fonds de la Recherche en Santé du Québec. SLA is an investigator of the Howard Hughes Medical Institute. The authors declare no competing financial interests.

## REFERENCES

Albrecht, S., Altenhein, B., and Paululat, A. (2011). The transmembrane receptor Uncoordinated5 (Unc5) is essential for heart lumen formation in Drosophila melanogaster. Dev Biol 350, 89–100.

Bhat, M.A., Rios, J.C., Lu, Y., Garcia-Fresco, G.P., Ching, W., St Martin, M., Li, J., Einheber, S., Chesler, M., Rosenbluth, J., et al. (2001). Axon-glia interactions and the domain organization of myelinated axons requires neurexin IV/Caspr/Paranodin. Neuron 30, 369–383.

Boyle, M.E., Berglund, E.O., Murai, K.K., Weber, L., Peles, E., and Ranscht, B. (2001). Contactin orchestrates assembly of the septate-like junctions at the paranode in myelinated peripheral nerve. Neuron 30, 385–397.

Bull, S.J., Bin, J.M., Beaumont, E., Boutet, A., Krimpenfort, P., Sadikot, A.F., and Kennedy, T.E. (2014). Progressive disorganization of paranodal junctions and compact myelin due to loss of DCC expression by oligodendrocytes. J Neurosci 34, 9768–9778.

Burgess, R.W., Jucius, T.J., and Ackerman, S.L. (2006). Motor axon guidance of the mammalian trochlear and phrenic nerves: dependence on the netrin receptor Unc5c and modifier loci. J Neurosci 26, 5756–5766.

Charles, P., Tait, S., Faivre-Sarrailh, C., Barbin, G., Gunn-Moore, F., Denisenko-Nehrbass, N., Guennoc, A.M., Girault, J.A., Brophy, P.J., and Lubetzki, C. (2002). Neurofascin is a glial receptor for the paranodin/Caspr-contactin axonal complex at the axoglial junction. Curr Biol 12, 217–220.

Chorghay, Z., Karadottir, R.T., and Ruthazer, E.S. (2018). White Matter Plasticity Keeps the Brain in Tune: Axons Conduct While Glia Wrap. Front Cell Neurosci 12, 428.

Cook, J.L., Irias-Donaghey, S., and Deininger, P.L. (1992). Regulation of rodent myelin proteolipid protein gene expression. Neurosci Lett 137, 56–60.

de Faria, O., Jr., Cui, Q.L., Bin, J.M., Bull, S.J., Kennedy, T.E., Bar-Or, A., Antel, J.P., Colman, D.R., and Dhaunchak, A.S. (2012). Regulation of miRNA 219 and miRNA Clusters 338 and 17-92 in Oligodendrocytes. Front Genet 3, 46.

de Faria, O., Jr., Pama, E.A.C., Evans, K., Luzhynskaya, A., and Karadottir, R.T. (2018). Neuroglial interactions underpinning myelin plasticity. Dev Neurobiol 78, 93–107.

Dhaunchak, A.S., Becker, C., Schulman, H., de Faria, O., Jr., Rajasekharan, S., Banwell, B., Colman, D.R., Bar-Or, A., and Canadian Pediatric Demyelinating Disease, G. (2012). Implication of perturbed axoglial apparatus in early pediatric multiple sclerosis. Ann Neurol 71, 601–613.

Dillon, A.K., Jevince, A.R., Hinck, L., Ackerman, S.L., Lu, X., Tessier-Lavigne, M., and Kaprielian, Z. (2007). UNC5C is required for spinal accessory motor neuron development. Mol Cell Neurosci 35, 482–489.

Downes, N., and Mullins, P. (2014). The development of myelin in the brain of the juvenile rat. Toxicol Pathol 42, 913–922.

Dupree, J.L., Coetzee, T., Blight, A., Suzuki, K., and Popko, B. (1998). Myelin galactolipids are essential for proper node of Ranvier formation in the CNS. J Neurosci 18, 1642–1649.

Einheber, S., Zanazzi, G., Ching, W., Scherer, S., Milner, T.A., Peles, E., and Salzer, J.L. (1997). The axonal membrane protein Caspr, a homologue of neurexin IV, is a component of the septate-like paranodal junctions that assemble during myelination. J Cell Biol 139, 1495–1506.

Fannon, A.M., Sherman, D.L., Ilyina-Gragerova, G., Brophy, P.J., Friedrich, V.L., Jr., and Colman, D.R. (1995). Novel E-cadherin-mediated adhesion in peripheral nerve: Schwann cell architecture is stabilized by autotypic adherens junctions. J Cell Biol 129, 189–202.

Fields, R.D. (2008). White matter in learning, cognition and psychiatric disorders. Trends Neurosci 31, 361–370.

Foran, D.R., and Peterson, A.C. (1992). Myelin acquisition in the central nervous system of the mouse revealed by an MBP-Lac Z transgene. J Neurosci 12, 4890–4897.

Freal, J.E., Kraft, G.H., and Coryell, J.K. (1984). Symptomatic fatigue in multiple sclerosis. Arch Phys Med Rehabil 65, 135–138.

Geisbrecht, B.V., Dowd, K.A., Barfield, R.W., Longo, P.A., and Leahy, D.J. (2003). Netrin binds discrete subdomains of DCC and UNC5 and mediates interactions between DCC and heparin. J Biol Chem 278, 32561–32568.

Glasgow, S.D., Labrecque, S., Beamish, I.V., Aufmkolk, S., Gibon, J., Han, D., Harris, S.N., Dufresne, P., Wiseman, P.W., McKinney, R.A., et al. (2018). Activity-Dependent Netrin-1 Secretion Drives Synaptic Insertion of GluA1-Containing AMPA Receptors in the Hippocampus. Cell Rep 25, 168–182 e166.

Glasgow, S.D., Ruthazer, E.S., and Kennedy, T.E. (2020a). Guiding synaptic plasticity: Novel roles for netrin-1 in synaptic plasticity and memory formation in the adult brain. J Physiol.

Glasgow, S.D., Wong, E.W., Thompson-Steckel, G., Marcal, N., Seguela, P., Ruthazer, E.S., and Kennedy, T.E. (2020b). Pre- and post-synaptic roles for DCC in memory consolidation in the adult mouse hippocampus. Mol Brain 13, 56.

Golan, N., Adamsky, K., Kartvelishvily, E., Brockschnieder, D., Mobius, W., Spiegel, I., Roth, A.D., Thomson, C.E., Rechavi, G., and Peles, E. (2008). Identification of Tmem10/Opalin as an oligodendrocyte enriched gene using expression profiling combined with genetic cell ablation. Glia 56, 1176–1186.

Goldman, J.S., Ashour, M.A., Magdesian, M.H., Tritsch, N.X., Harris, S.N., Christofi, N., Chemali, R., Stern, Y.E., Thompson-Steckel, G., Gris, P., et al. (2013). Netrin-1 promotes excitatory synaptogenesis between cortical neurons by initiating synapse assembly. J Neurosci 33, 17278–17289.

Gow, A., Southwood, C.M., Li, J.S., Pariali, M., Riordan, G.P., Brodie, S.E., Danias, J., Bronstein, J.M., Kachar, B., and Lazzarini, R.A. (1999). CNS myelin and sertoli cell tight junction strands are absent in Osp/claudin-11 null mice. Cell 99, 649–659.

Hamano, K., Takeya, T., Iwasaki, N., Nakayama, J., Ohto, T., and Okada, Y. (1998). A quantitative study of the progress of myelination in the rat central nervous system, using the immunohistochemical method for proteolipid protein. Brain Res Dev Brain Res 108, 287–293.

Heinemann, U., and Schuetz, A. (2019). Structural Features of Tight-Junction Proteins. Int J Mol Sci 20.

Hong, K., Hinck, L., Nishiyama, M., Poo, M.M., Tessier-Lavigne, M., and Stein, E. (1999). A ligand-gated association between cytoplasmic domains of UNC5 and DCC family receptors converts netrin-induced growth cone attraction to repulsion. Cell 97, 927–941.

Horn, K.E., Glasgow, S.D., Gobert, D., Bull, S.J., Luk, T., Girgis, J., Tremblay, M.E., McEachern, D., Bouchard, J.F., Haber, M., et al. (2013). DCC expression by neurons regulates synaptic plasticity in the adult brain. Cell Rep 3, 173–185.

Howell, O.W., Palser, A., Polito, A., Melrose, S., Zonta, B., Scheiermann, C., Vora, A.J., Brophy, P.J., and Reynolds, R. (2006). Disruption of neurofascin localization reveals early changes preceding demyelination and remyelination in multiple sclerosis. Brain 129, 3173–3185.

Jarjour, A.A., Bull, S.J., Almasieh, M., Rajasekharan, S., Baker, K.A., Mui, J., Antel, J.P., Di Polo, A., and Kennedy, T.E. (2008). Maintenance of axo-oligodendroglial paranodal junctions requires DCC and netrin-1. J Neurosci 28, 11003–11014.

Jarjour, A.A., Velichkova, A.N., Boyd, A., Lord, K.M., Torsney, C., Henderson, D.J., and Ffrench-Constant, C. (2020). The formation of paranodal spirals at the ends of CNS myelin sheaths requires the planar polarity protein Vangl2. Glia.

Kamasawa, N., Sik, A., Morita, M., Yasumura, T., Davidson, K.G., Nagy, J.I., and Rash, J.E. (2005). Connexin-47 and connexin-32 in gap junctions of oligodendrocyte somata, myelin sheaths, paranodal loops and Schmidt-Lanterman incisures: implications for ionic homeostasis and potassium siphoning. Neuroscience 136, 65–86.

Keleman, K., and Dickson, B.J. (2001). Short- and long-range repulsion by the Drosophila Unc5 netrin receptor. Neuron 32, 605–617.

Kennedy, T.E., Serafini, T., de la Torre, J.R., and Tessier-Lavigne, M. (1994). Netrins are diffusible chemotropic factors for commissural axons in the embryonic spinal cord. Cell 78, 425–435.

Kiernan, B.W., Garcion, E., Ferguson, J., Frost, E.E., Torres, E.M., Dunnett, S.B., Saga, Y., Aizawa, S., Faissner, A., Kaur, R., et al. (1999). Myelination and behaviour of tenascin-C null transgenic mice. Eur J Neurosci 11, 3082–3092.

Kosaras, B., and Kirschner, D.A. (1990). Radial component of CNS myelin: junctional subunit structure and supramolecular assembly. J Neurocytol 19, 187–199.

Krupp, L.B., Alvarez, L.A., LaRocca, N.G., and Scheinberg, L.C. (1988). Fatigue in multiple sclerosis. Arch Neurol 45, 435–437.

Lai Wing Sun, K., Correia, J.P., and Kennedy, T.E. (2011). Netrins: versatile extracellular cues with diverse functions. Development 138, 2153–2169.

Leonardo, E.D., Hinck, L., Masu, M., Keino-Masu, K., Ackerman, S.L., and Tessier-Lavigne, M. (1997). Vertebrate homologues of C. elegans UNC-5 are candidate netrin receptors. Nature 386, 833–838.

Lu, X., Le Noble, F., Yuan, L., Jiang, Q., de Lafarge, B., Sugiyama, D., Breant, C., Claes, F., de Smet, F., Thomas, J.L., et al. (2004). The netrin receptor UNC5B mediates guidance events controlling morphogenesis of the vascular system. Nature 432, 179–186.

Macabenta, F.D., Jensen, A.G., Cheng, Y.S., Kramer, J.J., and Kramer, S.G. (2013). Frazzled/DCC facilitates cardiac cell outgrowth and attachment during Drosophila dorsal vessel formation. Dev Biol 380, 233–242.

Macklin, W.B., Gardinier, M.V., Obeso, Z.O., King, K.D., and Wight, P.A. (1991). Mutations in the myelin proteolipid protein gene alter oligodendrocyte gene expression in jimpy and jimpymsd mice. J Neurochem 56, 163–171.

Maheras, K.J., Peppi, M., Ghoddoussi, F., Galloway, M.P., Perrine, S.A., and Gow, A. (2018). Absence of Claudin 11 in CNS Myelin Perturbs Behavior and Neurotransmitter Levels in Mice. Sci Rep 8, 3798.

Manitt, C., Colicos, M.A., Thompson, K.M., Rousselle, E., Peterson, A.C., and Kennedy, T.E. (2001). Widespread expression of netrin-1 by neurons and oligodendrocytes in the adult mammalian spinal cord. J Neurosci 21, 3911–3922.

Manitt, C., Thompson, K.M., and Kennedy, T.E. (2004). Developmental shift in expression of netrin receptors in the rat spinal cord: predominance of UNC-5 homologues in adulthood. J Neurosci Res 77, 690–700.

Marcus, J., Honigbaum, S., Shroff, S., Honke, K., Rosenbluth, J., and Dupree, J.L. (2006). Sulfatide is essential for the maintenance of CNS myelin and axon structure. Glia 53, 372–381.

Mathey, E.K., Derfuss, T., Storch, M.K., Williams, K.R., Hales, K., Woolley, D.R., Al-Hayani, A., Davies, S.N., Rasband, M.N., Olsson, T., et al. (2007). Neurofascin as a novel target for autoantibody-mediated axonal injury. J Exp Med 204, 2363–2372.

Miyamoto, T., Morita, K., Takemoto, D., Takeuchi, K., Kitano, Y., Miyakawa, T., Nakayama, K., Okamura, Y., Sasaki, H., Miyachi, Y., et al. (2005). Tight junctions in Schwann cells of peripheral myelinated axons: a lesson from claudin-19-deficient mice. J Cell Biol 169, 527–538.

Mombaerts, P., Wang, F., Dulac, C., Chao, S.K., Nemes, A., Mendelsohn, M., Edmondson, J., and Axel, R. (1996). Visualizing an olfactory sensory map. Cell 87, 675–686.

Newland, P., Starkweather, A., and Sorenson, M. (2016). Central fatigue in multiple sclerosis: a review of the literature. J Spinal Cord Med 39, 386–399.

Pedraza, L., Huang, J.K., and Colman, D.R. (2001). Organizing principles of the axoglial apparatus. Neuron 30, 335–344.

Podjaski, C., Alvarez, J.I., Bourbonniere, L., Larouche, S., Terouz, S., Bin, J.M., Lecuyer, M.A., Saint-Laurent, O., Larochelle, C., Darlington, P.J., et al. (2015). Netrin 1 regulates blood-brain barrier function and neuroinflammation. Brain 138, 1598–1612.

Poliak, S., Matlis, S., Ullmer, C., Scherer, S.S., and Peles, E. (2002). Distinct claudins and associated PDZ proteins form different autotypic tight junctions in myelinating Schwann cells. J Cell Biol 159, 361–372.

Schaeren-Wiemers, N., Bonnet, A., Erb, M., Erne, B., Bartsch, U., Kern, F., Mantei, N., Sherman, D., and Suter, U. (2004). The raft-associated protein MAL is required for maintenance of proper axon--glia interactions in the central nervous system. J Cell Biol 166, 731–742.

Scheiermann, C., Meda, P., Aurrand-Lions, M., Madani, R., Yiangou, Y., Coffey, P., Salt, T.E., Ducrest-Gay, D., Caille, D., Howell, O., et al. (2007). Expression and function of junctional adhesion molecule-C in myelinated peripheral nerves. Science 318, 1472–1475.

Schmittgen, T.D., Zakrajsek, B.A., Mills, A.G., Gorn, V., Singer, M.J., and Reed, M.W. (2000). Quantitative reverse transcription-polymerase chain reaction to study mRNA decay: comparison of endpoint and real-time methods. Anal Biochem 285, 194–204.

Schneider, C.A., Rasband, W.S., and Eliceiri, K.W. (2012). NIH Image to ImageJ: 25 years of image analysis. Nat Methods 9, 671–675.

Schuller, U., Heine, V.M., Mao, J., Kho, A.T., Dillon, A.K., Han, Y.G., Huillard, E., Sun, T., Ligon, A.H., Qian, Y., et al. (2008). Acquisition of granule neuron precursor identity is a critical determinant of progenitor cell competence to form Shh-induced medulloblastoma. Cancer Cell 14, 123–134.

Schultz, J., Milpetz, F., Bork, P., and Ponting, C.P. (1998). SMART, a simple modular architecture research tool: identification of signaling domains. Proc Natl Acad Sci U S A 95, 5857–5864.

Serafini, T., Kennedy, T.E., Galko, M.J., Mirzayan, C., Jessell, T.M., and Tessier-Lavigne, M. (1994). The netrins define a family of axon outgrowth-promoting proteins homologous to C. elegans UNC-6. Cell 78, 409–424.

Sherman, D.L., Tait, S., Melrose, S., Johnson, R., Zonta, B., Court, F.A., Macklin, W.B., Meek, S., Smith, A.J., Cottrell, D.F., et al. (2005). Neurofascins are required to establish axonal domains for saltatory conduction. Neuron 48, 737–742.

Simpson, S., Jr., Tan, H., Otahal, P., Taylor, B., Ponsonby, A.L., Lucas, R.M., Blizzard, L., Valery, P.C., Lechner-Scott, J., Shaw, C., et al. (2016). Anxiety, depression and fatigue at 5-year review following CNS demyelination. Acta Neurol Scand 134, 403–413.

Soriano, H.E., Kang, D.C., Finegold, M.J., Hicks, M.J., Wang, N.D., Harrison, W., and Darlington, G.J. (1998). Lack of C/EBP alpha gene expression results in increased DNA synthesis and an increased frequency of immortalization of freshly isolated mice [correction of rat] hepatocytes. Hepatology 27, 392–401.

Stevenson, B.R., and Keon, B.H. (1998). The tight junction: morphology to molecules. Annu Rev Cell Dev Biol 14, 89–109.

Tait, S., Gunn-Moore, F., Collinson, J.M., Huang, J., Lubetzki, C., Pedraza, L., Sherman, D. L., Colman, D.R., and Brophy, P.J. (2000). An oligodendrocyte cell adhesion molecule at the site of assembly of the paranodal axo-glial junction. J Cell Biol 150, 657–666.

Wallace, J.E., Krauter, E.E., and Campbell, B.A. (1980). Motor and reflexive behavior in the aging rat. J Gerontol 35, 364–370.

Winer, J., Jung, C.K., Shackel, I., and Williams, P.M. (1999). Development and validation of real-time quantitative reverse transcriptase-polymerase chain reaction for monitoring gene expression in cardiac myocytes in vitro. Anal Biochem 270, 41–49.

Wolswijk, G., and Balesar, R. (2003). Changes in the expression and localization of the paranodal protein Caspr on axons in chronic multiple sclerosis. Brain 126, 1638–1649.

Wong, E.W., Glasgow, S.D., Trigiani, L.J., Chitsaz, D., Rymar, V., Sadikot, A., Ruthazer, E. S., Hamel, E., and Kennedy, T.E. (2019). Spatial memory formation requires netrin-1 expression by neurons in the adult mammalian brain. Learn Mem 26, 77–83.

