## Supplemental figures 1-4 for "Paranode stability requires UNC5B expression by oligodendrocytes"

**Supplemental Figure 1**

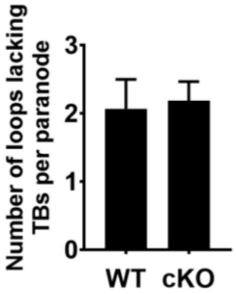


**Supplemental Figure 1: Lack of evidence for specific loss of transverse bands in UNC5B cKOs –** The mean number of glial loops that lack TBs per paranode was measured from EM micrographs from 7 month-old mutants and wild-types. 14-26 paranodes were analyzed. Bar graphs are plots of means and error bars indicate ±SEM.

**Supplemental Figure 2**

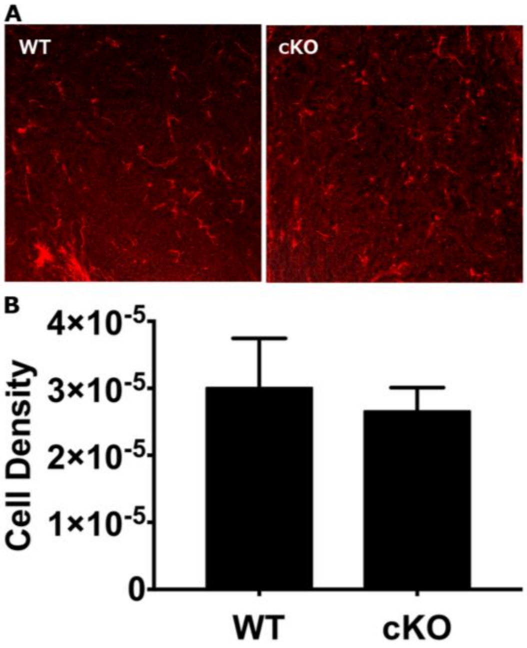


**Supplemental Figure 2: UNC5B cKOs display no signs of inflammation – (A)** 6-9 month-old UNC5B cKO and wild-type cerebellar sections were stained for the microglia marker CD31. **(B)** Quantification reveals no difference in microglia cell density. 4-5 animals were analyzed. Error bars indicate ±SEM.

**Supplemental Figure 3**

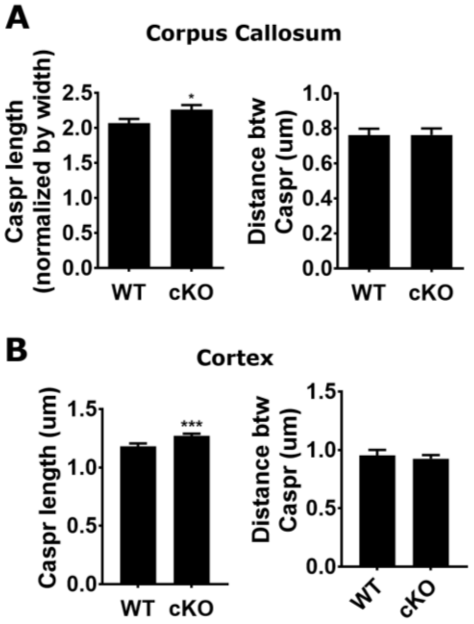


**Supplemental Figure 3: Caspr domain segregation is compromised in the corpus callosum and cortex of UNC5B cKO mice** – Measurements of Caspr1 domain length and distance between adjacent Caspr1 domains in the **(A)** corpus callosum and **(B)** cortex of 6-9 month old UNC5B cKO and wild-type controls. > 80 paranodes were measured. Error bars indicate ±SEM.

**Supplemental Figure 4**

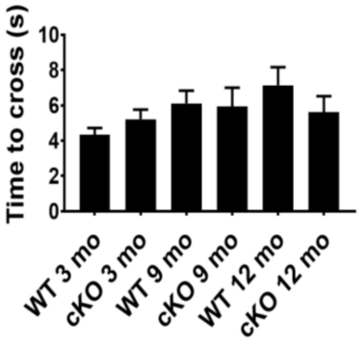


**Supplemental Figure 4: UNC5B cKO and WT littermates perform equally well in the beam walking test** – 6-14 mice for each genotype, at 3, 6-9 and 12 months, were trained to cross a 1 cm wide beam and the time to cross was measured. No significant differences were found between genotypes. Error bars indicate ±SEM.

**Supplemental Movie 1:** EM tomography of an UNC5B cKO paranode reveals fragmentation of the glial loop-loop interface that extends into the compact myelin.
